# High and low permeability of human pluripotent stem cell-derived Blood Brain barrier models depend on epithelial or endothelial features

**DOI:** 10.1101/2022.05.31.494120

**Authors:** Stéphane D. Girard, Ingrid Julien-Gau, Yves Molino, Benjamin F. Combes, Louise Greetham, Michel Khrestchatisky, Emmanuel Nivet

## Abstract

The search for reliable human blood-brain barrier (BBB) models represents a challenge for the development/testing of strategies aiming to enhance brain delivery of drugs. Human induced pluripotent stem cells (hiPSCs) have raised hopes in the development of predictive BBB models. Differentiating strategies are thus required to generate endothelial cells (ECs), a major component of the BBB. Several hiPSC-based protocols have reported the generation of *in vitro* models with significant differences in barrier properties. We studied in depth the properties of iPSCs byproducts from two protocols that have been established to yield these *in vitro* barrier models. Our analysis/study reveals that iPSCs endowed with EC features yield high permeability models, while the cells that exhibit outstanding barrier properties show principally epithelial cell-like (EpC) features. Our study demonstrates that hiPSC-based BBB models need extensive characterization beforehand and that a reliable human BBB model is still needed.

## INTRODUCTION

The blood vessels that vascularize the central nervous system (CNS) possess unique properties and are collectively referred to as the blood brain barrier (BBB). The BBB is composed of specialized brain microvascular endothelial cells (BMVECs) that establish intercellular tight junctions, surrounded by pericytes, astrocytes and neurons, altogether forming the neurovascular unit (NVU). CNS vessels are non-fenestrated and BMVECs tightly regulate CNS homeostasis, notably the movement of ions, molecules, and cells between the blood and the brain (Abbott et al., 2010). These properties rely in part on the expression of BBB-specific receptors, transporters and efflux pumps at the level of BMVECs. Interactions of BMVECs with other cells of the NVU precisely control the brain microenvironment and protect the CNS from toxic compounds, pathogens, inflammation, injury, and disease (Aday et al., 2016, Larsen et al., 2014). BBB dysfunctions and inflammation accompany neurodegenerative disorders (NDDs) indicating that it plays a key role in these disorders (Sweeney et al., 2018). If the highly selective permeability of the BBB is essential to preserve CNS integrity from a large variety of putative toxic products, it also represents a major obstacle to deliver pharmaco-active compounds to the brain (Patel and Patel, 2017).

Accordingly, the use of robust *in vitro* BBB models recapitulating at least in part phenotypical and functional characteristics is of major relevance to study BBB physiopathology as well as to predict the cerebral exposure of new neuro-pharmaceuticals (Wolff et al., 2015). *In vitro* BBB models produced from animal brain microvessels have shown to be reliable and very useful for studying the BBB (Helms et al., 2016). However, the establishment of predictive human cell-based BBB models is still required to address inter-species differences at the molecular, cellular and functional levels. Until recently, modeling of the human BBB was restricted to the use of human primary BMVECs or to immortalized human cell lines, both presenting limitations for drug screening and trans-endothelial transport evaluation. For instance, primary human BMVECs usually obtained from post-mortem tissue or patient biopsies have obvious limitations regarding availability, scalability and reproducibility, and human cell lines have failed to show optimal barrier properties (Aday et al., 2016, Helms et al., 2016).

Recently, new promising strategies based on the use of other cellular sources such as human cord blood-derived hematopoietic stem cells or endothelial progenitors have been proposed to produce brain-like endothelial cells (Boyer-Di Ponio et al., 2014, Cecchelli et al., 2014). Concomitantly, the use of human induced pluripotent stem cells (hiPSCs) has gained large interest as starting material for alternative strategies (Lauschke et al., 2017, Ferreira, 2019, Appelt-Menzel et al., 2020). Indeed, in addition to their human origin, these cells present several advantages including unlimited cell source (scalability) and the potential to differentiate into any cell types (versatility) and, in the BBB context, to generate the different cellular components of the NVU (Lippmann et al., 2013). Moreover, hiPSCs are easily available and can be produced from patient cells to study genetic diseases and potentially BBB dysfunctions associated with specific diseases/mutations (Bosworth et al., 2017, Logan et al., 2019).

A prerequisite for using hiPSCs as BBB modeling tools is the development of efficient protocols to direct their differentiation into functional BMVECs. On the one hand, several studies, based on chemically defined methods, have described the conversion of pluripotent stem cells (PSCs) into endothelial cells (ECs) with variable efficiencies (Kurian et al., 2013, Lian et al., 2014, Patsch et al., 2015, Liu et al., 2016, Nguyen et al., 2016, Lin et al., 2017, Belt et al., 2018, Halaidych et al., 2018, Olmer et al., 2018, Farkas et al., 2020, Nishihara et al., 2021). These procedures are based on the use of small molecules and growth factors activating key signaling pathways aiming to recapitulate early developmental stages of EC generation. Similar differentiation paradigms, based on three main steps, are generally described: a mesodermal induction followed by an endothelial/vascular induction that precedes an EC expansion phase coupled with a cell sorting procedure to obtain nearly pure EC populations. However, to our knowledge, the attempts to specify these ECs into BMVEC-like cells have not yet been fully satisfactory and these cells show insufficient barrier properties (Minami et al., 2015, Lauschke et al., 2017, Praca et al., 2019).

On the other hand, Lippmann and collaborators demonstrated the possibility to obtain cells with exceptional barrier properties from human PSCs using a fundamentally different procedure (Lippmann et al., 2012). This strategy, which was subsequently improved (Lippmann et al., 2014, Wilson et al., 2015, Wilson et al., 2016, Hollmann et al., 2017, Qian et al., 2017, Neal et al., 2019) and adapted by many different groups (Katt et al., 2016, Mantle et al., 2016, Appelt-Menzel et al., 2017, Kokubu et al., 2017, Kurosawa et al., 2018, Ribecco-Lutkiewicz et al., 2018, Ohshima et al., 2019, Roux et al., 2019, Neal et al., 2021), is based on an undirected/spontaneous differentiation method that is supposed to promotes neural and endothelial co-differentiation. These cells have also been used in complex microfluidic systems (DeStefano et al., 2017, Wang et al., 2017, Grifno et al., 2019, Motallebnejad et al., 2019, Park et al., 2019, Vatine et al., 2019, Jagadeesan et al., 2020) and combined with other NVU cells derived from the same hiPSC lines to produce isogenic human BBB models (Canfield et al., 2017, Canfield et al., 2019, Blanchard et al., 2020). The generation of patient-derived cells using this undirected method to study BBB dysfunction in neurological disorders has also been reported (Lim et al., 2017, Vatine et al., 2017, Lee et al., 2018, Oikari et al., 2020).

In the search of identifying the best available strategy to establish a reliable human BBB modeling platform for further applications, we performed a thorough analysis of the cellular phenotype and barrier properties generated by undirected/spontaneous differentiation of hiPSCs (Lippmann et al., 2014, Lippmann et al., 2012). Results were compared to the cellular phenotype and barrier properties of hiPSC differentiated using an alternative protocol based on a chemically defined mesodermal induction (Patsch et al., 2015), but also of primary BMVECs. Our study shows that chemically defined mesodermal induction of hiPSCs generated cells with endothelial-like features, but rather poor barrier properties. In contrast, undirected/spontaneous differentiation of hiPSCs yielded exquisite barrier properties but with cells that presented essentially an epithelial-like phenotype.

## RESULTS

### The undirected hiPSC-based differentiation protocol produces cells lacking expression of critical EC markers

To seek for the best method able to reproducibly generate hiPSC-derivatives suitable for BBB modeling, we first characterized differentiated cells from three independent hiPSC lines (see supplementary table 1) by comparing two published protocols (Fig. 1A). The first protocol (hereafter referred as Pr-1) was based on a 5-day induction phase with a chemically defined method followed by cell sorting to purify ECs prior to expansion (Challet Meylan et al., 2015, Patsch et al., 2015). The second protocol (hereafter referred as Pr-2) relied on an 8-day non-chemically defined differentiating method (i.e. resembling embryoid bodies spontaneous differentiating protocols) and further cell purification by sub-culturing them onto a collagen IV/fibronectin matrix (Lippmann et al., 2014, Stebbins et al., 2016). By day 11 post-differentiation, both strategies led to the generation of cells that formed a uniform monolayer after seeding at a defined density on a collagen IV/fibronectin coated membrane (Fig. 1B). However, the differentiated cells exhibited significant morphological differences (Fig. 1B). Cells obtained with Pr-1 showed a typical elongated spindle-shaped morphology while cells derived with Pr-2 displayed cobblestone-like morphology. Of note, as recommended by Pr-1, the cells were initially expanded in a culture medium containing VEGF that is an important angiogenic factor well known to induce BBB disruption and, consequently, not compatible for BBB modeling. Thus, for further experiments, we decided to remove VEGF after seeding onto filters. Nevertheless, we noticed that the cell monolayer obtained with Pr-1 derivatives was strongly disorganized upon VEGF removal (Supplementary Fig. 1). To bypass this issue, we established a specific seeding density (0.5 x 10^6^ cells/cm^2^) accompanied by a gradual decrease of the VEGF concentration in the culture media that allowed maintaining the organization of the cell monolayer obtained with Pr-1 (Fig. 1B).

**Figure 1.**
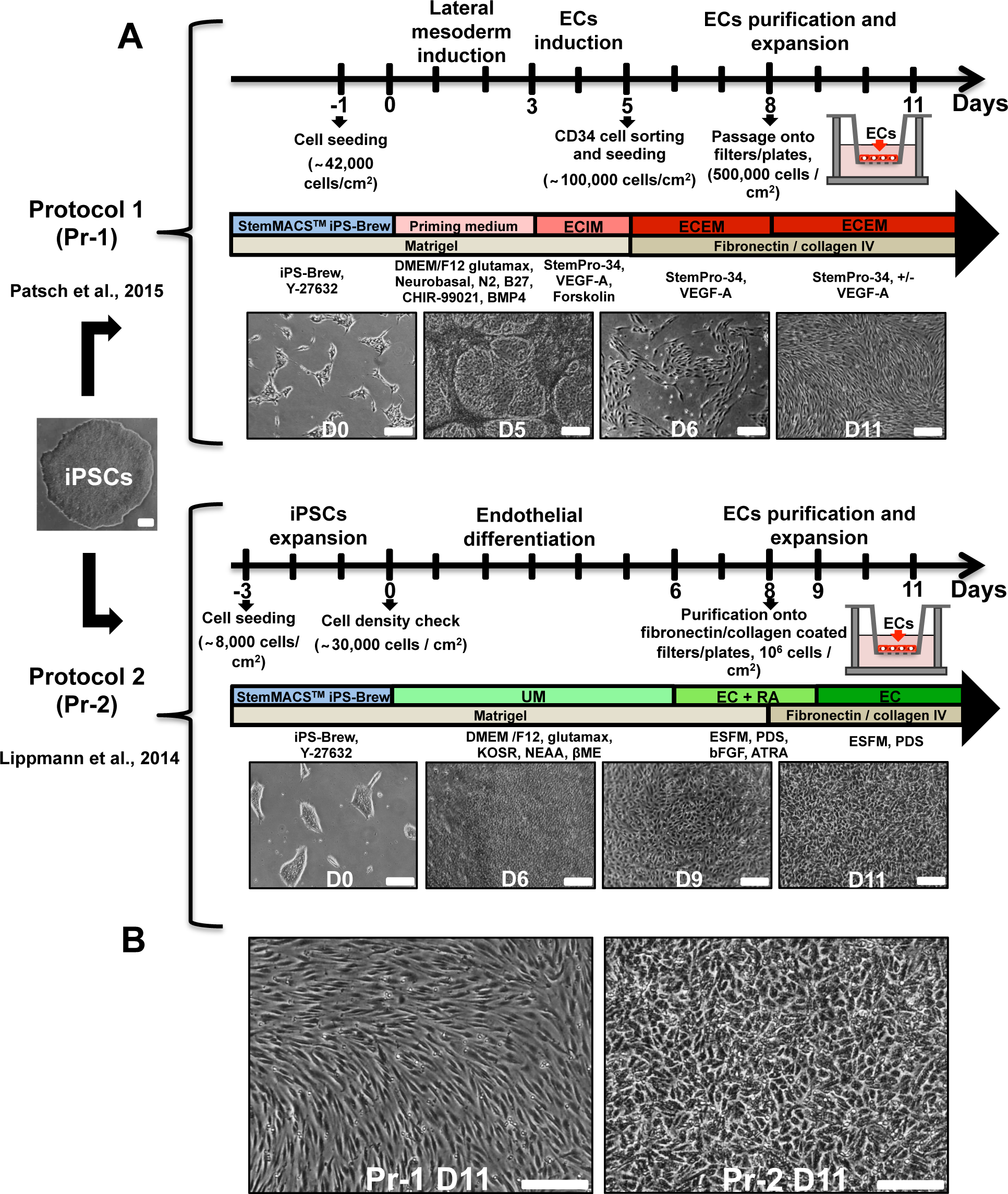
Representative overview of the two hiPSCs-based differentiation protocols used in this study. **A)** Timelines summarizing the different steps of the two differentiation protocols used throughout the study, namely Pr-1 (top, adapted from Patsch et al., 2015) and Pr-2 (bottom, adapted from Lippmann et al., 2014). A summary of the main components of the different culture media and coatings used at each steps of the protocols is indicated and representative bright-field micrographs illustrate the morphology of the cells at different differentiation stages where D stands for Days of differentiation. **B)** Higher magnification micrographs corresponding to cells representative of cell population obtained upon completion of the two differentiation protocols and seeded on filters, at day 11 (D11). Scale bars, 0.2 mm (A, B).

The strong morphological differences obtained with the two protocols led us to question whether the cells generated from Pr-1 and Pr-2, respectively, shared similar expression of EC markers. We first assessed the expression of three cell surface markers, namely CD31/PECAM1, CD34 and CD144/VE-Cadherin. For both protocols, the dynamic expression of these EC markers was followed by flow cytometry at different stages of the differentiation: undifferentiated state (i.e. hiPSC stage or day 0), before sorting or purification (day 5 or 8) and after expansion of the sorted/purified cells (day 11). HUVECs (peripheral ECs) and primary human BMVECs (HBMVECs) were used as positive control for EC markers while the Caco-2 cell line, representative of epithelial cells (EpCs), was used as negative control (Supplementary Fig. 2). On all three independent hiPSC lines we tested, the Pr-1 led to a high yield of differentiating cells displaying strong expression of all three markers, even before purification (Fig. 2A, 2B and Supplementary Fig. 2A). Thus, before sorting, a high percentage of hiPS-derivatives obtained from Pr-1 were found positive for CD31 (77.5 +/- 7.4%) and CD34 (94.3 +/- 2%). After cell sorting, nearly pure populations of cells expressing CD31, CD34 and CD144 were isolated by day 11. Differentiated cells generated with Pr-2 showed no CD31 expression before and after purification while 97.5 +/-0.1% of the primary HBMVECs were positive for this major endothelial marker (Fig. 2A, 2B and Supplementary Fig. 3A). Furthermore, CD34 expression level in the Pr-2 differentiated cells was significantly lower compared to all three undifferentiated hiPSC lines (Fig. 2A and 2B). To assess whether the extracellular matrix (ECM) could impact the differentiation efficacy, we tested two different ECM concentrations of collagen type IV/ fibronectin: i) a mixture of 10 µg/mL of both collagen type IV and fibronectin, according to our own expertise using primary rat BMEC (Molino et al., 2014) (Mixt. 1) and ii) a mixture of 400 µg/ml collagen type IV and 100 µg/ml fibronectin, based on the Pr-2 recommendation (Mixt.2). In both situations, the Pr-2 protocol failed to generate CD31+ and CD34+ cells (Supplementary Fig. 3B). We next ruled out that the absence of CD31+ cells upon Pr-2-based differentiation was due to the anti-CD31 antibody we used as two distinct antibodies were tested and showed similar results (Supplementary Fig. 3C). Moreover, we also tested whether the media used to maintain and amplify hiPSCs could impact on the fate of those undifferentiated cells upon differentiating conditions. Thus, we tested the Pr-2 protocol on hiPSCs maintained in either mTeSR^TM^1 or iPS-BREW and no difference was observed, confirming that the Pr-2 protocol did not generate CD31+ cells and produced very low numbers of CD34+ cells (Supplementary Fig. 3D). Noteworthy, out of the three EC markers we preselected and analyzed, only CD144 was induced by Pr-2, compared to undifferentiated hiPSCs (Fig. 2A and 2B). Moreover, after the purification procedure, 33% of CD144 + cells were obtained with the Pr-2 protocol at most. These results were further confirmed by immunocytochemistry analyses that revealed a weak and highly heterogeneous CD144 signal at the intercellular junctions of Pr-2 cells in comparison with Pr-1 cells and HBMVECs (Fig. 2C). Similar differences were also observed with the Von Willebrand Factor (VWF), another EC marker that was strongly expressed within Pr-1 cells and HBMVECs and poorly detected in Pr-2 cells. Based on these observations, we next decided to deepen the characterization of the Pr-2 cells by assessing their functional ability to uptake fluorescent acetylated LDL, a characteristic of endothelial cells (Voyta et al., 1984). Flow cytometry analyses revealed that sorted CD34+ Pr-1 cells were able to uptake Alexa 488 Ac-LDL similarly to HUVECs and HBMVECs while purified Pr-2 cells displayed a very weak signal as for fibroblasts that were used as negative control (Fig. 2A-C). Matrigel capillary-like tube formation assay was additionally performed, confirming that only cells generated with Pr-1 were able to form tube-like structures similar to vascular cells such as HBMVEC or HUVECs (Fig. 2D and Supplementary Fig. 3E). Of note, we confirmed that the inability of Pr-2 derived cells to generate HBMVEC-like tubes was not due to cell viability prior to seeding (Supplementary Fig. 3E).

**Figure 2.**
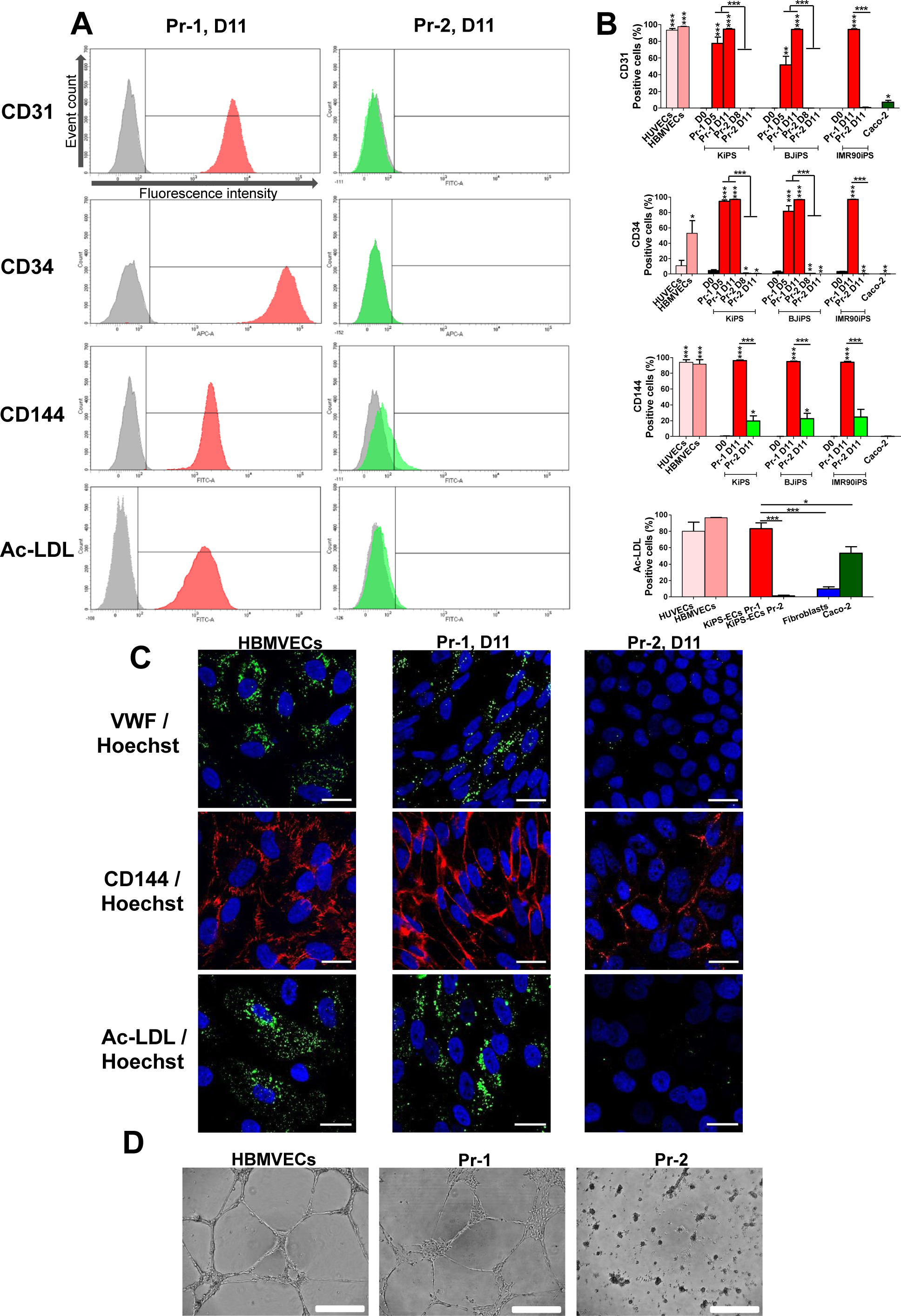
Pr-2-derived cells fail to express classical hallmarks of ECs. **A)** Representative flow cytometry results showing fluorescence intensity of stained cells in comparison with appropriate isotype controls (grey) in differentiated cells according to Pr-1 (red) and Pr-2 (green) protocols at day 11 (D11). The analyses revealed the absence of classical EC surface markers (as indicated) in Pr-2-derived cells, contrary to Pr-1-derived cells. Pr-2-derivatives were also found unable to uptake fluorescent acetylated LDL (Ac-LDL). **B)** Quantification of the percentage of positive cells for the different EC markers and Ac-LDL in hiPSC-derived cells (three independent hiPSC lines) at different stages of the two protocols (D5 and D11) and in cells used as positive (HUVECs and HBMVEC) and negative (Caco-2) endothelial controls. Values reported are the mean (± SEM) of at least three independent differentiations/cultures with *p ≤ 0.05, **p ≤ 0.01, and ***p ≤ 0.001, using Student’s t-test (if not specified, comparison with respective D0 undifferentiated control). **C)** Representative fluorescent micrographs showing immunostainings of EC markers including VWF (green, top row), CD144 (red, middle row) and acetylated LDL (green, bottom row). Nuclei were counterstained with Hoechst (blue). **D)** Representative bright-field micrographs of the indicated cells 24h after Matrigel capillary-like tube formation assay. Scale bars: 20 µm (C) and 500 µm (D).

Our observations led us to question whether the cells generated by Pr-2 expressed other EC markers or whether their remarkable barrier properties (Lippmann et al., 2014) were related to an epithelial phenotype. We thus analyzed by RT-qPCR the gene expression levels of a defined set of 9 EC markers (*PECAM1*, *CD34*, *CDH5*, *VWF*, *ENG*, *FLT1*, *KDR*, *TIE1* and *TEK*) and 4 EpC markers (*CDH1*, *EPCAM*, *KRT8* and *CLDN4*) and compared with those of ECs (HUVECs, HBMVECs and HCMEC/D3) and EpCs (Caco-2) used as controls. Confirming our protein expression data, we observed that EC markers were strongly induced in CD34+ Pr-1 cells when compared to their undifferentiated counterparts (Fig. 3A and Supplementary Fig. 4). The gene expression profile of all 9 markers analyzed from Pr-1 cells was similar in HUVECs, HBMVECs and HCMEC/D3, further confirming their EC-like identity. In contrast, purified Pr-2 cells expressed EC markers at low or similar levels when compared with undifferentiated hiPSCs, with the exception of *CDH5*/*CD144* as previously observed at the protein level. Noteworthy, our analyses also revealed that the expression of the 4 EpC markers in Pr-2 cells (Fig. 3B) was similar to that of undifferentiated hiPSCs, and reminiscent of the Caco-2 gene expression profile. Conversely, these 4 EpC markers were under-expressed in Pr-1 cells, HUVECs, HBMVECs and HCMEC/D3 in comparison with Caco-2 (Fig. 3B). For all genes analyzed (Fig. 3A-B), significant differences were observed between cells obtained with these two protocols (p < 0.05).

**Figure 3.**
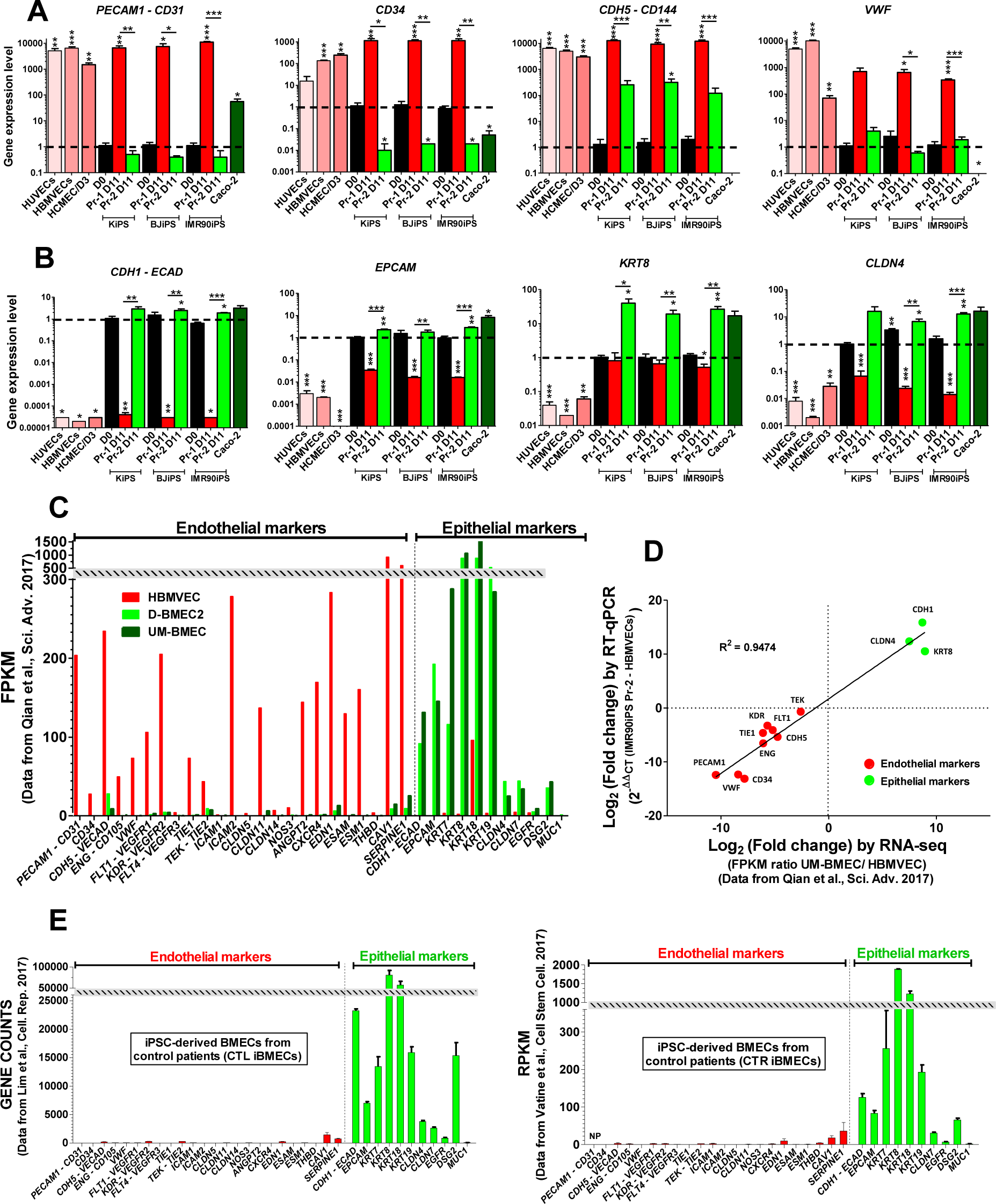
Transcriptomic analyses revealed that Pr-2-derived cells display an epithelial-like phenotype. Bar graphs showing the mRNA expression profile of a defined set of EC markers **(A)** and EpC markers **(B)** assessed by RT-qPCR in hiPSC-derived cells at different stages of the two protocols. EC (HUVECs, HBMVECs, HCMEC/D3) and EpC (Caco-2) cells were used as positive controls. Values reported are shown as a fold change relative to the value of undifferentiated iPSCs (D0) and are represented as a mean (± SEM) of at least three independent differentiations/cultures with *p ≤ 0.05, **p ≤ 0.01, and ***p ≤ 0.001, using Student’s t-test. **C)** Bar graphs showing the gene expression profile of two sets of cell markers representative of endothelial or epithelial cells, respectively. Gene expression levels were from RNAseq analyses obtained by a previous study (Qian et al., 2017) in HBMVECs and in D10 IMR90iPS-derived cells differentiated according to a “Pr-2-like” protocol (UM-BMEC) or to a chemically defined method (D-BMEC2). Values reported are FPKM extracted from GEO database (accession number GSE97575). **D)** Correlative analysis between the relative gene expression of the selected set of genes as measured in the present study by RT-qPCR and in the previous study by RNAseq (Qian et al., 2017). The value represents the log2 of the fold change between HBMVECs and IMR90iPS-derived ECs differentiated according to the Pr-2 protocol (RT-qPCR) or a “Pr-2-like” protocol (UM-BMEC, RNAseq). **E)** Two additional examples of RNAseq data extracted from GEO database showing the gene expression profiles of EC and EpC markers in other hiPSC-derived ECs differentiated according to a “Pr-2-like” protocols (Left, raw read count abundance; Right, reads per kilobase million (RPKM) accession numbers: GSE97100 (Lim et al., 2017) and GSE97324 (Vatine et al., 2017)). NP: not published.

It is not likely that our observations result from differences between the cell lines used in our study and those of Lippmann and colleagues, considering that we included in our study the IMR90-4 hiPSC line, which was originally used to set up Pr-2 (Lippmann et al., 2014, Lippmann et al., 2012). Altogether, our results suggested that cells generated by Pr-2 failed to express characteristic EC markers, at both genic and proteic levels. Moreover, those Pr-2 cells lacked critical EC features such as capillary tube formation. Taken together, our data suggest that Pr-1 and Pr-2 generate cells with EC- and EpC-like phenotypes, respectively.

### Pr-2-derived cells display an EpC-like phenotype

We further questioned whether our observations resulted from in house experimental bias or from Pr-2. To this end, we leveraged on transcriptomic data sets generated by other groups using Pr-2 and compared them with our gene expression analyzes. We extracted RNA sequencing data (RNA-Seq) from the Gene Expression Omnibus (GEO) database. We selected two groups of genes (24 EC and 10 EpC genes) and directly reported the values uploaded by different authors on the database. We started by analyzing the data previously generated from HBMVECs and purified IMR90-4iPS-derived cells (Qian et al., 2017) according to a “Pr-2-like” protocol (UM-BMVEC) and to a variant procedure (D-BMVEC2) described by the authors as a more chemically defined method allowing production of brain ECs in a directed manner (GEO accession number GSE97575). Our analyses revealed that HBMVECs and “Pr-2-like” cells derived from hiPSCs displayed highly different gene expression profiles (Fig. 3C). HBMVECs expressed all EC and EpC genes at high and low levels respectively when compared to UM-BMVEC and D-BMVEC2. Noticeably, critical EC markers such as *PECAM1*, *CD34* and *CLDN5* displayed zero or near zero fragment per kilobase million (FPKM) values in “Pr-2-like” cells. Importantly, the D-BMVEC2 cells derived from the “chemically defined method” exhibited the same gene expression profile. We also compared the fold change values (Pr-2-derived cells *versus* HBMVECs) obtained from our RT-qPCR study (Fig. 3A-B) with those obtained by RNA-seq reported by Qian and collaborators (Qian et al., 2017), and showed a high correlation as indicated by the linear regression analysis (r^2^ = 0.9474; Fig. 3D). Furthermore, we also found these same expression profiles – i.e. EC and EpC – from RNAseq data available from four other studies based on “Pr-2-like” protocols (Fig. 3E and Supplementary Fig. 5C). In all cases, the reported data indicated a lack or a very low expression of EC genes in comparison with EpC genes (GEO accession numbers: GSE97100 (Lim et al., 2017), GSE97324 (Vatine et al., 2017), GSE108012 (Lee et al., 2018) and GSE129290 (Vatine et al., 2019)). To support our findings, we also compared these results with other data sets from RNAseq studies performed in human nasal and intestinal EpCs (GEO accession numbers: GSE 107898 (Landry and Foxman, 2018) and GSE94935 (Lickwar et al., 2017)). Here again, we found an expression profile comparable to Pr-2 cells (Supplementary Fig. 5B) unlike that we observed in others ECs (Supplementary Fig. 5A) such as HUVECs and mouse brain microvascular ECs (GEO accession numbers: GSE93511 (Zhang et al., 2017) and GSE111839 (Sabbagh et al., 2018)).

Finally, we performed further comparison of the Pr-1 and Pr-2 generated cells by studying the expression of EpC protein markers using flow cytometry analyses. We showed that only Pr-1-derived cells lose the expression of Ep-CAM and E-cadherin across the differentiation process, whereas Pr-2-derived cells strongly expressed these two epithelial markers throughout differentiation at levels similar to those of Caco-2 cells (Supplementary Fig. 6A-B). Complementary immunocytochemical analyses against cytokerain-8 and cytokeratin-18, two other epithelial markers, confirmed their strong expression in undifferentiated hiPSC, Pr-2-derived cells and Caco-2 cells whereas these markers were barely detectable in Pr-1-derived cells (Supplementary Fig 6C). Altogether, analyses at the transcriptomic and protein levels provide further evidence indicating that Pr-2 cells display an epithelial-rather than endothelial-like phenotype.

### Pr-2-derived EpCs express BBB-specific markers

Pr-2 cells have been largely used for BBB modeling and we next sought to deepen their characterization. We first analyzed the gene expression levels of a pool of 8 BBB markers including 3 tight junction related genes (*TJP1-ZO1*, *CLDN5*, *OCLN*) and 5 genes coding for BBB transporters and efflux pumps (*ABCB1-PGP*, *SLC2A1-GLUT1*, *TFRC*, *LDLR*, *SLC7A5-LAT1*). As above, CD34-sorted Pr-1 cells and purified Pr-2 cells were compared to undifferentiated hiPSCs, peripheral and brain ECs (HUVECs, HBMVECs and HCMEC/D3) as well as to epithelial cells (Caco-2). For most of the selected genes, significant differences (p < 0.05) between Pr-1 and Pr-2 cells were found with greater expression of BBB markers in Pr-2-derived cells. This was true for essential BBB genes such as *OCLN*, *PGP* and *GLUT1*, with the exception of *CLDN5* that was found strongly overexpressed in Pr-1 cells and in the other ECs but nearly absent in Pr-2-derived cells (Fig. 4A).

**Figure 4.**
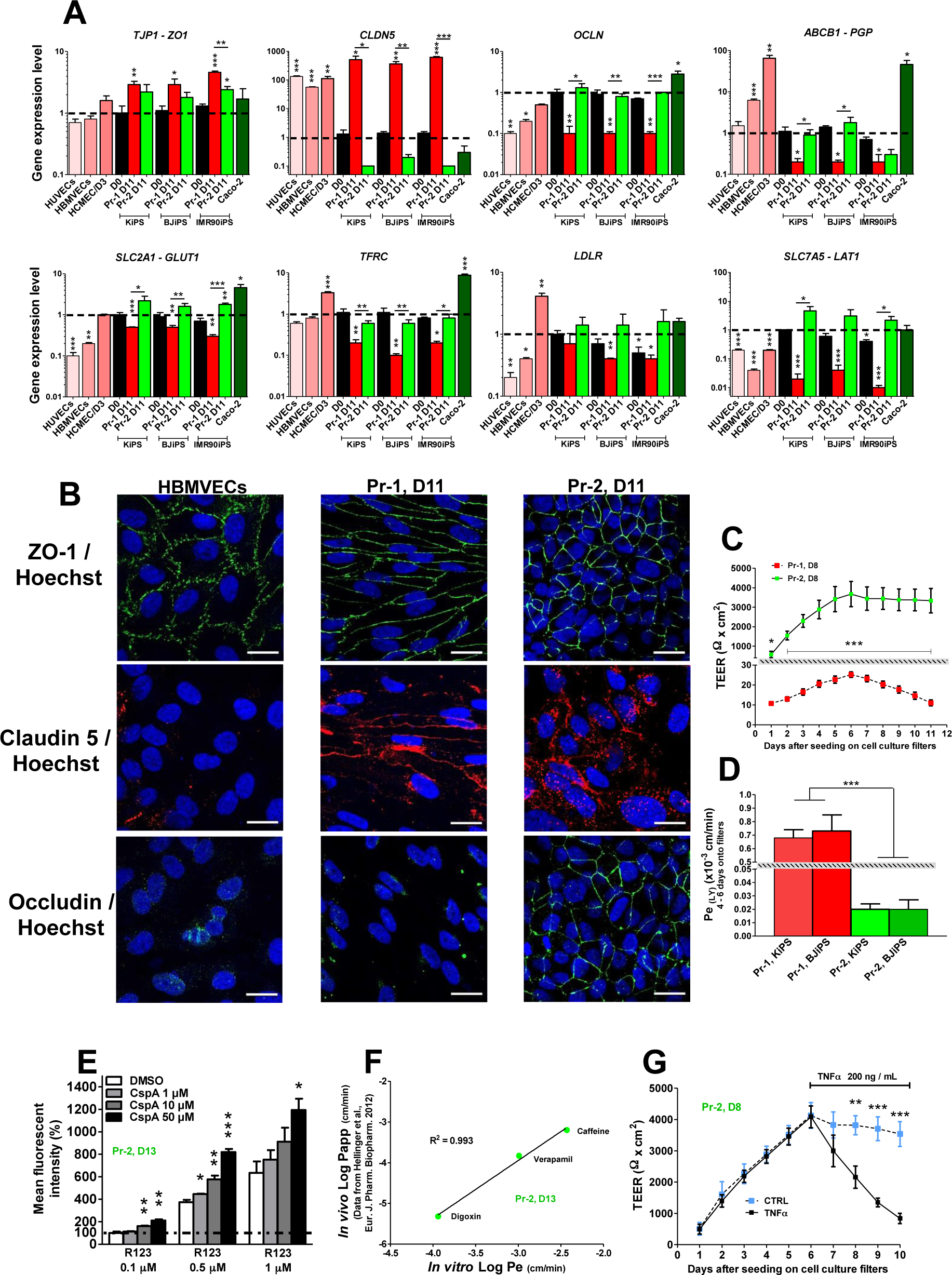
Evaluation of BBB markers and barrier properties from Pr-1 and Pr-2 derived cells. **A)** Bar graphs showing the mRNA expression profile of the indicated BBB markers as assessed by RT-qPCR in hiPSC-derived cells at day 11 of differentiation with either Pr-1 or Pr-2. EC (HUVECs, HBMVECs, HCMEC/D3) and EpC (Caco-2) were used as controls. Values reported are shown as fold change relative to the value of undifferentiated hiPSCs (D0) and are displayed as a mean (± SEM). **B)** Representative fluorescent micrographs showing immunostainings against tight junction proteins including ZO-1 (green, top row), Claudin 5 (red, middle row) and Occludin (green, bottom row). Nuclei were counterstained with Hoechst (blue). **C)** Graph showing kinetics of TEER results in Pr-1 and Pr-2-derived KiPS cells over a period of 11 days after seeding as monolayers on filters at D8 of differentiation. **D)** The paracellular permeability to lucifer yellow (LY) was assessed from Pr-1 and Pr-2-derived KiPS and BJiPS cells 4 to 6 days after seeding on filters at D8 of differentiation. Results obtained from cells derived from two independent hiPSC lines are shown. **E)** P-glycoprotein (P-GP) efflux activity was measured by intracellular accumulation of Rhodamine 123 (R123) using flow cytometry in the absence (DMSO, vehicle) or presence of the P-GP specific inhibitor cyclosporin A (CspA) in Pr-2-derived ECs. **F)** Correlation between *in vitro* permeability coefficients (Pe, x-axis) obtained in KiPS-derived Pr-2 ECs cultured as monolayers and the *in vivo* apparent permeability coefficients (Papp) previously measured in mice brains ((Hellinger et al., 2012) of three different drugs as indicated. **G)** Graph showing kinetics of TEER in Pr-2 derivatives in the presence or absence of TNF-α and showing that TNF-α treatment altered the functional integrity of the barrier properties. For all experiments, the values reported are representative of at least three independent differentiations/cultures with *p ≤ 0.05, **p ≤ 0.01, and ***p ≤ 0.001 using Student’s t-test. Scale bars: 20 µm.

We also assessed the expression of three classical markers for tight junctions (Z0-1, Claudin-5 and Occludin) by immunolabeling of cell monolayers obtained either with Pr-1, Pr-2 or primary HBMVECs (Fig. 4B). Although ZO-1 expression was clearly detectable at the intercellular junctions in all three monolayers, significant differences were observed for Claudin-5 and Occludin. Claudin-5 expression was heterogeneously distributed at the junctions between Pr-1-derived cells, appeared disorganized in Pr-2-derived monolayers and was barely detectable in cultures obtained with primary HBMVECs. Occludin was only detectable at the intercellular junctions in Pr-2-derived monolayer cultures. Altogether, our observations indicated that Pr-2-derived EpCs better recapitulated the expression of BBB markers readily related to barrier tightness when compared to Pr-1-derived ECs but also primary HBMVECs.

### Pr-2-derived EpCs display strong barriers properties

Based on the aforementioned observations, we then performed analyses at the functional level to evaluate the tightness of the above-mentioned cell monolayers by assessing the trans-endothelial electrical resistance (TEER) and the paracellular permeability to lucifer yellow (LY) (Fig. 4C-D). After cell seeding on the top of a microporous membrane classically used for BBB modeling, we performed a dynamic analysis for 11 days of the TEER of cultures derived from sorted Pr-1 cells and purified Pr-2 cells, which revealed strong significant differences from the very first day (p < 0.001). After 5 days of seeding Pr-2 cells reached a very high TEER value (3676.5 +/- 644 Ω.cm^2^) that lasted for at least seven days whereas the TEER values obtained with the Pr-1 cells remained below 30 Ω.cm^2^ (Fig. 4C). Similarly, significant differences were obtained when we analyzed LY permeability (Fig. 4D). Pr-1 cells reached a mean Pe (LY) of 0.7 x 10^-3^ +/- 0.1 cm/min while Pr-2 cells exhibited a mean Pe (LY) of 0.02 x 10^-3^ +/- 0.007 cm/min after 5 days on filters (p < 0.001). Of note, it is classically considered that TEER values of about 150-200 Ω.cm^2^ are the lowest functional limit for *in vitro* models (Reichel et al., 2003, Wolff et al., 2015). Consequently, in view of the low tightness measured with Pr-1-derived cells that we could not be optimized, we only pursued the functional characterization of the Pr-2 cells and their monolayers. We used flow cytometry analysis to measure the functionality of the P-glycoprotein (P-GP) efflux pump by assessing the intracellular accumulation of Rhodamine 123 (R123), a cell permeable P-GP substrate, in absence or presence of cyclosporin A (CspA), a P-GP specific inhibitor (Fig. 4E). We observed a dose-dependent effect of the inhibitor demonstrated by an increase of the R123 accumulation in purified Pr-2 cells, which started to be significant at 1 µM of CspA with 0.5 µM of R123 (p<0.05) when compared to the vehicle control. We observed a 2.05-fold increase (+/- 0.1) in R123 cellular accumulation in the presence of 50 µM CspA. In addition, we assessed the *in vitro* permeability (Pe) through Pr-2 cell monolayers of three small molecules (Digoxin, Verapamil and Caffeine) known to display different *in vivo* brain penetration because of their different size, lipophilicity, and ability to bind to efflux transporters (Hellinger et al., 2012). The *in vitro* Pe values of these molecules measured by LC/MS1MS and *in vivo* apparent permeability coefficients (Papp) previously measured in mice brains (Hellinger et al., 2012) showed a very high correlation as indicated by the linear regression analysis (r^2^ = 0.993) (Fig. 4F). We completed the characterization of the Pr-2 monolayers by analyzing the effect of a pro-inflammatory cytokine (TNFα) on their integrity (Fig. 4G). The TEER kinetic study indicated a strong decrease of the tightness of cytokine-treated monolayers, which started to be significant after 48h of treatment (p<0.001). Overall, different functional assays confirmed that Pr-2-derived cells displayed strong barrier properties.

## DISCUSSION

Human iPSCs have enormous potential for the development of human *in vitro* models that can provide new platforms for fundamental research, but also for screening/testing/validation of molecules in the context of preclinical studies on humanized systems. As a prerequisite, such models need to closely recapitulate both phenotypic and functional features of the tissue(s) or cell type(s) to be studied. In this perspective, establishing a reliable and scalable human BBB model is paramount for the study of active molecules on the human BBB, their ability to cross this physical barrier, but also to develop strategies to deliver molecules that cannot naturally pass the BBB. Considering that BMVECs represent one of the key components of the BBB, one major challenge for BBB modeling is the generation of hiPSC-derived ECs displaying a cerebral phenotype and that reproduce *in vitro* the landmarks of the BBB *in vivo*.

### Pr-2-derived cells form a model with good barrier properties but show limited endothelial features

In 2012, the Shusta laboratory (Lippmann et al., 2012) reported an efficient method to produce human cells endowed with characteristics of the *in vivo* BBB. The cells produced with this method, which was subsequently improved in a series of follow-up reports (Lippmann et al., 2014, Stebbins et al., 2016, Qian et al., 2017), have been described as ECs exhibiting good barrier properties with TEER values up to 8,000 Ω.cm^2^ (Blanchard et al., 2020) exceeding by far TEERs usually observed with primary BMVECs from different species (Wolff et al., 2015, Helms et al., 2016). The proposed method is relatively simple as it does not require multiple steps involving the application of well-defined titers of small molecules and growth factors known to activate key developmental signaling pathways, which is usually the strategy followed to differentiate PSCs into ECs (Patsch et al., 2015, Kurian et al., 2013, Nguyen et al., 2016). Indeed, during the first 6 days of this differentiation procedure, small hiPSC colonies solely require DMEM and knock-out serum, indicating that removal of factors that maintain the pluripotent state of hiPSCs, is sufficient to produce differentiated cells efficiently. The only molecules added to the differentiation medium the next three days (day 6 to day 9) are bFGF in the first report (Lippmann et al., 2012) and, according to following reports, all-trans retinoic acid that significantly improved the barrier properties (Lippmann et al., 2014). Moreover, this method involves purification of cells by a sub-culture onto collagen IV/fibronectin matrix and does not require cell sorting procedures based on endothelial surface markers (such as CD31, CD34 or CD144), as usually performed to obtain nearly pure EC populations from PSCs (Patsch et al., 2015, Kurian et al., 2013, Nguyen et al., 2016, Lian et al., 2014). These authors reported that their procedure is based on PSCs co-differentiation into both neural and endothelial lineages providing an “embryonic brain-like microenvironment” allowing the specification of ECs into BMVECs *via* a Wnt/β-catenin signaling (Lippmann et al., 2012). Although it is well accepted that the embryonic brain environment is key for BMVECs specification, the latter have a mesodermal origin distinct from the neuro-ectodermal neural cells and acquire their BBB properties only after migrating into the developing brain from the perineural vascular plexus (Engelhardt, 2003, Obermeier et al., 2013, Kurz, 2009). Most studies describing the generation of ECs from PSCs include a defined mesodermal induction step based on the use of specific molecules such as glycogen synthase kinase 3 β (GSK3β) inhibitors, BMP4 or Activin A. This mesoderm induction step is usually combined with activators of the endothelial phenotype such as VEGF, cyclic AMP activators, or inhibitors of transforming growth factor beta (TGFβ) signaling (Liu et al., 2016, Patsch et al., 2015, Kurian et al., 2013, Belt et al., 2018, Nguyen et al., 2016, Lian et al., 2014, Lin et al., 2017, Minami et al., 2015). In the present study, we compared cells obtained in the absence of such a mesodermal-induction step (Pr-2) with cells produced with a fully defined procedure (Pr-1). Our results showed that Pr-2 promotes the generation of epithelial-like cells rather than ECs, as demonstrated by transcriptomic and proteomic analyses. Some reports have already pointed out drawbacks regarding the phenotype of cells derived from the protocol of Lippmann and collaborators. For instance, Delsing et al. reported that these cells may exhibit a mixed endothelial and epithelial phenotype while Vatine and collaborators indicated that they share similarities with epithelial cells (Delsing et al., 2018, Vatine et al., 2019). Another study mainly based on single-cell RNA sequencing and bioinformatic analysis reported that the protocol of Lippmann and collaborators generates EPCAM^+^/PECAM1^-^ neuro-ectodermal epithelial cells lacking endothelial identity (Lu et al., 2021). In addition, it has been shown that Pr-2 lack expression of several key adhesion molecules making these cells not suitable to study the interaction of immune cells with the BBB (Nishihara et al., 2020). These reports recently led Lippmann and collaborators to rename these cells as “BMEC-like cells” and to indicate that Pr-2 cells are not identical to human BMECs *in vivo* and therefore may not be appropriate for all applications (Lippmann et al., 2020). Nevertheless, these cells are still referred to as brain endothelial cells. Two observations appear to support this appellation.

First, like others, we also showed that Pr-2-derived cells expressed key BBB markers including tight junction proteins such as ZO-1 and Occludin but also efflux pump transporters. However, with the exception of the loss of Claudin 5 expression in Pr-2-EpCs, we noted that BBB-specific markers were already expressed in undifferentiated hiPSCs with no major changes upon differentiation with Pr-2. Hence, our observations are in phase with previous reports showing that undifferentiated iPSCs exhibit an epithelial phenotype resulting from a mesenchymal-to-epithelial transition during the reprogramming process (Li et al., 2010, Teshigawara et al., 2017). This seems to indicate that some key BBB features observed in Pr-2-derived cells and associated with an epithelial phenotype are not necessarily induced by Pr-2 but rather maintained during their differentiation.

Second, our data confirm the original and following reports showing that cells obtained with the Pr-2 method displayed solid barrier properties at the functional level that probably explain why this method has been used extensively. In particular, different reports using this differentiation protocol demonstrated good correlations between *in vitro* and *in vivo* barrier permeability of standard molecules known to display different *in vivo* brain penetration (Lippmann et al., 2012, Mantle et al., 2016, Roux et al., 2019, Ohshima et al., 2019, Ribecco-Lutkiewicz et al., 2018). This has led several groups to use this model as a tool to predict barrier permeability of small compounds or larger biomolecules with therapeutic interest (Clark et al., 2016, Gallagher et al., 2016, Chiou et al., 2018, Mantle and Lee, 2019). Although the Pr-2 model generate cells endowed with EpC rather than EC features, it may be relevant to test molecules that cross barriers *via* cellular mechanisms shared by BMVECs and EpCs. However, it may not be appropriate for mechanistic studies on the transport of molecules across HBMVECs or to study other mechanisms such as immune cell interactions at the BBB (Nishihara et al., 2020). This comparison led to unexpected results concerning the phenotype of the Pr-2 cells, raising concerns regarding their use for hiPSC-based BBB models.

### Pr-1-derived cells show EC features but form barrier models with low TEER

We compared in our study the differentiation and barrier properties of hPSCs generated by Pr-2 and Pr-1. The latter is based on a chemically defined method with mesodermal induction that is conceptually optimal from a developmental perspective. We selected the protocol published by Patsch et al., that describes the efficient and rapid conversion of PSCs into ECs with an efficiency exceeding 80% within six days of differentiation (Patsch et al., 2015). The procedure (Challet Meylan et al., 2015) was easily reproduced in our hands with two slight modifications. First, we replaced the mTSeR^TM^1 medium used to culture the undifferentiated hiPSCs by the StemMACS iPS-Brew medium. Second, at the end of the differentiation process, we replaced the CD144 antibody by a CD34 antibody to improve the yield of magnetic cell sorting (not shown), a strategy that has already been used in similar differentiation protocols (Minami et al., 2015, Lian et al., 2014, Kurian et al., 2013). As expected, we produced from three different hiPSCs lines pure EC populations with high efficiency and reproducibility. Contrary to the Pr-2 procedure, Pr-1-derived cells displayed a full EC phenotype including the expression of all major EC markers, formed vascular tubes *in vitro* and demonstrated uptake of acetylated LDL similarly to primary ECs (HBMVECs and HUVECs) used as controls. In comparison with HUVECs, Pr-1-derived ECs displayed higher CD34 expression (both at the protein and mRNA levels) but this marker is known to be downregulated in primary HUVECs after several passages (Fina et al., 1990). Also, even though we did not verify this point in the present study, it has been shown that Pr-1-derived ECs form vascular structures *in vivo* after transplantation which validates their angiogenic potential (Patsch et al., 2015). To our knowledge, this ability has never been demonstrated for Pr-2-derived cells and recently Lu and collaborators reported that these cells do not form lumenized vessels in immunocompromised mice (Lu et al., 2021).

However, the Pr-1 procedure does not appear to generate ECs with brain specialization (BMVECs). Thus, although we obtained homogeneous monolayers of contiguous cells (Fig. 1 and 4), we showed that Pr-1 cells exhibit poor barrier properties including low TEER values. These values were close to those previously described for other hiPSCs-derived ECs produced with other chemically defined strategies, indicating that these hiPSC-ECs are close to peripheral ECs displaying low TEER such as HUVECs (Patsch et al., 2015, Minami et al., 2015, Lian et al., 2014). We also observed no or weak expression of key BBB markers such as Occludin and P-GP.

In order to improve the barrier properties of these “unspecified / naïve” cells, we tested different strategies aiming at reproducing a cerebral environment based either on co-cultures with cerebral cells (neural progenitors, glial cells, brain pericytes) or on cultures with molecules known to promote BMVEC specification: angiopoietin-1, Wnt-3a, TGFβ1, sonic hedgehog, retinoic acid, adrenomedullin and cyclic AMP activators. However, there was no significant improvement of Pr-1 barrier properties (not shown). As a consequence, we considered that in our experimental conditions, the cells generated using Pr-1 are not suitable for transport experiments of small or large molecules. Overall, our results were consistent with previous studies based on chemically defined methods that showed insufficient barrier properties (Lauschke et al., 2017, Minami et al., 2015, Praca et al., 2019, Nishihara et al., 2020) considering that a relevant *in vitro* BBB model should exhibit TEER values of at least 150-200 Ω.cm^2^ for transport experiments (Reichel et al., 2003, Wolff et al., 2015).

New differentiating methods are thus needed to specify hiPSC-derived ECs or hiPSC-derived vascular progenitors (Kurian et al., 2013) towards a BMVEC phenotype optimal for BBB modeling. This will probably require a better understanding of the cellular and molecular mechanisms, including signaling pathways that occur during the specification of ECs in the developing brain. Recently, by comparing the transcriptomic profile of non- and CNS-derived murine ECs (Roudnicky et al., 2020a) or by performing a compound library screening on a CLDN5-GFP PSC line (Roudnicky et al., 2020b), Roudnicky and collaborators identified new factors that control barrier resistance in ECs and showed a moderate improvement of the barrier properties in Pr-1-derived cells. Considering that the cerebral microenvironment made of and controlled by NVU cells is probably key to BMVEC specification and maintenance (Engelhardt, 2003, Obermeier et al., 2013), another way to improve the cerebral phenotype of hiPSC-derived ECs could be to more closely mimic the complexity of the cerebrovascular interface *in vitro*. Considerable efforts are currently being made to develop “mini brain” with 3D cytoarchitecture such as brain organoids, spheroids or “brain-on-a-chip” whose interest goes beyond the simple modeling of the BBB (Bhalerao et al., 2020). Human iPS cells by their ability to differentiate into different NVU cells have an important role to play in the development of these new innovative technologies. Recently, brain organoids produced from hiPSCs have been vascularized by hiPSC-derived ECs generated with a chemically-defined procedure close to the Pr-1 protocol (Pham et al., 2018). Other strategies such as the supplementation of VEGF in the differentiation medium (Ham et al., 2020) or the ectopic induction of the expression of specific ETS transcription factors (Cakir et al., 2019) have been used to induce the co-differentiation of ECs and the vascularization of brain organoids. Further studies will determine whether these new technological developments will not only help to better understand the neurovascular interactions but will also allow the establishment of reliable human BBB models (Waldau, 2019).

In conclusion, using different approaches and relevant cellular controls, we demonstrated that cells obtained with the Pr-2 differentiation procedure require further characterization. They display an epithelial rather than endothelial phenotype and our observations are supported by transcriptomic data from several studies. However, cells derived from this procedure may be of great value in some cases, especially due to their barrier properties. As an alternative, cells generated by the Pr-1 procedure display EC features but lack the full differentiation into BMVECs yielding *in vitro* models with poor barrier properties.

## EXPERIMENTAL PROCEDURES

### Cell culture

Three human iPSC lines were used as previously characterized: KiPS 4F2 (Aasen et al., 2008), BJIPS 6F (Xia et al., 2013) and iPS (IMR90)-4 (Yu et al., 2007) (Supplementary table 1). Human iPSCs (hiPSCs) were cultured in a chemically defined growth medium (StemMACS iPS-Brew XF (Miltenyi Biotec) or mTSeR^TM^1 (Stem Cell Technologies) on plates coated with growth factor reduced matrigel (8.7 µg/cm^2^, Corning). For passaging, 70-80% confluent iPSCs were treated with a cell dissociation buffer (0.5 mM EDTA, 1.8 mg/mL NaCl, D-PBS without Ca^2+^/Mg^2+^) for 3 min at RT, and colonies were dispersed to small clusters using a 5-ml glass pipette and carefully replated at a splitting ratio of about 1:4. Expression of pluripotent stem cell transcription factors (Nanog, Oct3/4, and Sox2) by the three human iPSC lines was controlled by flow cytometry, using BD Stemflow Human Pluripotent Stem Cell Transcription Factor Analysis Kit® (BD Biosciences) according to the manufacturer’s instructions. Human umbilical vein endothelial cells (HUVECs) were cultured in EBM-2 medium supplemented with EGM-2 SingleQuots (Lonza) on plates coated with rat tail collagen I (20 µg/cm^2^, Corning). Human brain microvascular endothelial cells (HBMVECs) were cultured in Endothelial Growth Medium (EGM, Angio-proteomie, PELOBiotech) on plates coated with human fibronectin (10 µg/mL, Corning). The hCMEC/D3 cells were cultured in EBM-2 medium supplemented with EGM-2 MV SingleQuots (Lonza) on plates coated with rat tail collagen I (20 µg/cm^2^, Corning). For RT-qPCR experiments, hCMEC/D3 cells grown at confluence were differentiated for 5 days in the same medium but without the growth factors. Caco-2 and human skin fibroblast cells were grown in Dulbecco’s modified Eagles medium (DMEM) with Glutamax (Life Technologies) and supplemented with 10% fetal bovine serum, 100 units/mL of penicillin and 100 µg/mL of streptomycin (Life Technologies). All cell types were maintained at 37°C in a humidified incubator at 5% CO2 with medium changes every day (hiPSCs) or every second day for the other cell types.

### Differentiation of hiPSCs into ECs

#### Protocol 1 (Pr-1)

This differentiation procedure is based on the protocol reported by Patsch et al. (Challet Meylan et al., 2015, Patsch et al., 2015) with slight modifications. In short, hiPSCs at 70-80% of confluence were dissociated with StemPro Accutase (Life technologies) and singularized cells were seeded at a defined density (between 42,000 and 62,000 cells/cm^2^ depending on the hiPSC line) on matrigel coated 6-well plates in StemMACS iPS-Brew supplemented with 10 µM Y27632 (ROCK inhibitor, Tocris Bioscience). After 24 h (Fig. 1), the medium was replaced with Priming Medium (1:1 mixture of DMEM/F12 with Glutamax and Neurobasal media, supplemented with 1x N2 and B27 (Life Technologies), 8 µM CHIR99021 (Tocris Bioscience), 25 µg/ml human recombinant Bone Morphogenetic Protein 4 (BMP4, R&D Systems) and 55 µM β-mercaptoethanol (Life technologies). After three days in the medium, it was replaced for two days by EC Induction Medium (ECIM: StemPro-34 SFM medium, Life Technologies) supplemented with 200 ng/ml human recombinant Vascular Endothelial Growth Factor (VEGF165, PeproTech) and 2 µM forskolin (Sigma-Aldrich). The ECIM was renewed the following day. On day five ECs were dissociated with StemPro Accutase (diluted 1:6 in D-PBS) and separated via Magnetic-activated cell sorting (MACS, Miltenyi Biotec) using CD34 MicroBeads, LS Columns and MidiMACS Separator according to the manufacturer’s protocol. CD34 positive cells were replated on dishes coated with 10 µg/mL human fibronectin (Corning) at a density of 1 x 10^5^ cells/cm^2^ in EC Expansion Medium (ECEM: StemPro-34 SFM with 100 ng/ml VEGF). ECEM was replaced every other day. Once confluence was reached (usually three days), the cells were either cryopreserved or directly seeded in ECEM (with 50 ng/ml VEGF) medium at a cell density of 0.5 x 10^6^ cells/cm^2^ on microporous (1 µm) polyethylene membrane (12- or 6-well insert filters, Greiner Bio-One) to establish EC monolayers or at a density of 0.25 x 10^6^ cells/cm^2^ on tissue culture-treated plates for other experiments. Filters and plates were coated (24h, 37°C) with a mixture of collagen type IV and fibronectin (10 µg/mL for both, Corning) as we previously published for rat brain microvascular cells (Molino et al., 2014). The next day, the medium was changed with ECEM (with or without 50 ng/ml VEGF) and then changed every other day.

#### Protocol 2 (Pr-2)

This differentiation procedure is based on the protocol reported by Lippmann et al. (Lippmann et al., 2014, Lippmann et al., 2012, Wilson et al., 2015, Stebbins et al., 2016). HiPSCs grown at 70-80% of confluence were dissociated with StemPro Accutase and singularized cells were seeded at a defined density (between 7,500 and 15,000 cells/cm^2^ depending on the hiPSC line) on Matrigel coated 6-well plates in StemMACS iPS-Brew or mTSeR^TM^1 media supplemented for the first 24h with 10 µM Y27632 (Fig. 1). Three days later, once the cells had reached an optimal cell density of about 3 x 10^4^ +/- 5 x 10^3^ cells/cm^2^, differentiation was initiated and medium changed to an “unconditioned medium” for 6 days with daily changes (UM: DMEM/F-12 + glutamax, 20% KnockOut Serum Replacement, 1x non-essential amino acids and 0,1 mM β-mercaptoethanol (Life technologies)). On day six, UM was changed for 2 days to an endothelial cell medium supplemented with retinoic acid (EC+RA: human Endothelial-SFM (Life Technologies), 1% platelet-poor plasma derived bovine serum (Alfa Aesar), 20 ng/mL human recombinant basic fibroblast growth factor (bFGF, PeproTech) and 10 µM all-trans retinoic acid (Sigma-Aldrich). On day eight, cells were passaged using StemPro Accutase (about 30 min at 37°C) and then either cryopreserved (Wilson et al., 2016) or directly subcultured in EC-RA medium at a cell density of 1 x 10^6^ cells/cm^2^ on microporous membrane to establish EC monolayers (see Pr-1) or at a density of 0.5 x 10^6^ cells/cm^2^ on culture-treated polystyrene plates for other experiments. Filters and plates were precoated (24h, 37°C) with a mixture of 400 µg/ml collagen type IV and 100 µg/ml fibronectin (Corning) as previously reported by Lippmann et al. (Lippmann et al., 2014) or with the same mixture at 10 µg/mL (Molino et al., 2014). The next day the medium was changed with EC medium without bFGF and retinoic acid, and then changed every other day.

### Flow cytometry analysis

Cells were washed twice in D-PBS and then harvested using StemPro Accutase. Cells were then centrifuged (300 x g, 5 min) and washed once with cold blocking solution (10% FBS in D-PBS). Expression of EC surface markers (CD31, CD34 and CD144) were performed on living cells. Cells were incubated with the corresponding antibodies or appropriate isotype controls (supplementary table 2) in cold blocking solution for 1h on ice in the absence of light. After incubation, cells were washed thrice with cold blocking solution and resuspended in a total volume of 200 µl before analysis. Acquisitions were performed on a FACSCanto II flow cytometer (BD Biosciences) using BDFACSDiva software. At least 10,000 events were recorded for each analysis and measures were performed in duplicate. Percentages of cells or mean fluorescence intensity are presented after the subtraction of isotype background and refer to the total population analyzed. Results are representative of at least three independent experiments with a minimum of two technical replicates per experiment.

In order to exclude bias due to potential enzymatic cleavages of the proteins of interest (surface antigens, EC or EpC markers) during the cell detachment procedure, the same enzymatic solution (same duration and same concentration) was applied on the different cell types for comparative purposes.

### Immunocytochemistry and fluorescence microscopy

Cells grown on filters were washed thrice with D-PBS and fixed in 4% paraformaldehyde (PFA, Antigenfix, MM France) for 15 min. Filters with the monolayers were gently dissociated from the plastic inserts with a razor blade before immunocytochemistry. Cells were then blocked and permeabilized for 30 min at RT in blocking buffer that contained 3% bovine serum albumin (BSA) and 0.1% Triton X-100 (Sigma Aldrich). Subsequently, cells were incubated 90 min at RT with the indicated primary antibody (supplementary table 2) diluted in PBS with 3% BSA. Cells were then washed thrice with PBS and incubated for 1h RT with the appropriate secondary antibodies and Hoechst 33342 (1 µg/mL Sigma-Aldrich). Cells were washed thrice with PBS and membranes mounted in Prolong Gold antifade mounting medium (Thermo Fisher Scientific). Negative control conditions were carried out by omitting the primary antibody. Images were acquired and processed using a confocal microscope (LSM 700) and Zen software (Carl Zeiss).

### Acetylated Low Density Lipoprotein (Ac-LDL) uptake assay

For fluorescence microscopy experiments, cells grown at confluence on filters were incubated with 10 µg/mL Alexa Fluor™ 488 Ac-LDL (Thermo Fisher Scientific) in DMEM/F-12 supplemented with 1% BSA for 4h at 37°C. The cells were washed thrice with PBS and nuclei counterstained with Hoechst 33342 before uptake assessment using confocal microscopy. For flow cytometry analysis, the cells grown at confluence on 24-well plates were incubated with 1 µg/mL Alexa Fluor™ 488 Ac-LDL for 30 min at 37°C. The cells were washed thrice with D-PBS, dissociated using TrypLE Select (Life technologies) and analyzed by flow cytometry.

### Vascular tube formation assay

Briefly, 96-well culture plates were coated with 50 µL/well of growth factor reduced matrigel (undiluted, about 1.25 mg/cm^2^) for 1h at 37°C. Cells were dissociated using StemPro Accutase and 100 µl of a cell suspension was dispensed in each well in their respective culture media supplemented with 50 ng/mL VEGF. For each cell type, different cell densities were tested (from 20 x 10^3^ to 140 x 10^3^ cells/cm^2^). Tube formation was observed and imaged after 24h of incubation.

### RNA isolation and real-time quantitative PCR analysis

Total RNA was isolated using the RNeasy Plus Micro Kit (Qiagen) according to the manufacturer’s recommendations. RNA concentration and purity were determined using a NanoDrop-1000 Spectrophotometer (NanoDrop Technologies, Thermo Fisher Scientific). According to the manufacturer’s protocol (Thermo Fisher Scientific), 0.5 µg total RNA was submitted to reverse transcription using Moloney Murine Leukemia Virus (MMLV) reverse transcriptase to generate single-strand cDNA. RT-qPCR experiments were performed with the 7500 Fast Real-Time PCR System (Applied Biosystems/Life Technologies). All reactions were performed using primers listed in the supplementary table 3 and the iTaq Universal SYBR Green Supermix (Bio-Rad Laboratories). Relative expression levels were determined according to the ΔΔCt method with the human housekeeping gene *ABELSON* as endogenous control for normalization as previously described (Beillard et al., 2003). For each condition, RNA extractions were performed from three independent cultures and the reported values are the mean fold change relative to the value of the control sample (undifferentiated cell line).

### Trans-endothelial electrical resistance (TEER) assay

Trans-endothelial electrical resistance (TEER) measurements were performed using a Millicell-ERS (Millipore) volt-ohmmeter connected to an EndOhm chamber (World Precision Instruments) suited for 12-well inserts. TEER measurements were performed in culture medium 24h after cell seeding on filters and every 24h until completion of the kinetic experiment (*i.e*. 10 days), according to the manufacturer’s recommendations. The resistance value (Ω.cm^2^) of an empty filter coated with collagen/fibronectin were used as blanks and subtracted from each measurement. All TEER measurements were performed in triplicates (at least three inserts per condition). TEER measurements were also performed on cell monolayers treated with Tumor Necrosis Factor-α (TNF-α). In this case, both upper and lower compartment were treated with 200 ng/mL TNF-α (PeproTech) and TEER measurements performed 24 h later and every 24 h for a 10 day-period. TNF-α treatment was renewed every other day.

### Permeability experiments

The monolayer tightness in 12-well inserts was also assessed by measuring LY (Lucifer Yellow CH dilithium salt, Sigma-Aldrich) permeability as previously described (Molino et al., 2014). Briefly, LY was incubated in the upper compartment of the culture system in contact with ECs for 60 min at 37°C in culture medium. Quantification of the LY paracellular leakage was assessed by fluorimetric analysis (excitation at 430 nm and emission at 535 nm) and expressed in LY permeability, Pe (LY).

The *in vitro* permeability of three substances, Digoxin, Verapamil and caffeine (Sigma-Aldrich, resuspended in DMSO), was assessed across the EC monolayers. Briefly, these three compounds were incubated at 4 µM in culture medium for 240 min at 37°C in the upper compartment of HTS 96-well Transwell inserts (Corning). The monolayer integrity was controlled using LY at the end of the experiment. The concentrations of the molecules in samples from upper and lower compartments were quantified using a validated liquid chromatography with tandem mass spectrometry (LC/MS1MS) method at Eurofins/ADME Bioanalyses (Vergeze, France). For each molecule, 12 inserts were quantified. Permeability (Pe) was calculated for each drug as previously described for LY (Molino et al., 2014).

### Functional assay for P-glycoprotein

P-GP efflux activity was assessed by analyzing the intracellular accumulation of rhodamine 123 (R123), a P-GP substrate incubated with or without the addition of the P-GP inhibitor cyclosporin A (CspA). Briefly, the cells grown at confluence on 24-well plates were pre-incubated with inhibitor at (1, 10 or 50 µM) or with vehicle (DMSO, Dimethyl sulfoxide) in DMEM/F-12 supplemented with 1% BSA for 30 min at 37°C. Then the cells were incubated with R123 (0.1, 0.5 or 1 µM) with or without inhibitor in the same medium for 1h at 37°C. The cells were then washed once in culture medium and incubated for 90 min at 37°C with or without inhibitor. The cells were washed thrice with D-PBS, dissociated using TrypLE Select and analyzed by flow cytometry. Duplicate samples were measured. Assays were performed at least 3 times.

### Statistics

All data are presented as mean ± SEM of at least three independent differentiations/cultures as indicated in the figure captions. Values were compared using Student’s t test. The minimal threshold for significance was set at p < 0.05.

## AUTHOR CONTRIBUTIONS

S.D.G designed and performed experiments, analyzed data, and drafted the manuscript. I.J.G, Y.M and B.C helped to perform experiments. L.G was involved in cell culture procedures. M.K and E.N led the conception, supervised the project and wrote the manuscript.

## ACKNOWLEDGEMENTS

We thank Françoise Jabès for her technical support regarding the *in vitro* BBB experiments and Guillaume Jacquot for his help concerning the analysis of small molecules permeability. This work was in part supported by funding from the Centre National de la Recherche Scientifique (CNRS) and Aix-Marseille Université, and by a public grant from the French National Research Agency (ANR, project “NANOVECTOR”, ANR-15-CE18-0010). This work has received support from the French government under the Programme Investissements d’Avenir, Initiative d’Excellence d’Aix-Marseille Université via A*Midex (AMX-19-IET-004) and ANR (ANR-17-EURE-0029) funding.

## CONFLICT of INTEREST

M.K. is director of the UMR7051 Laboratory, and cofounder, shareholder, and scientific counsel of the Vect-Horus biotechnology company. The remaining authors declare no conflicts of interest.

## FIGURE LEGENDS

**Supplementary Figure 1.**
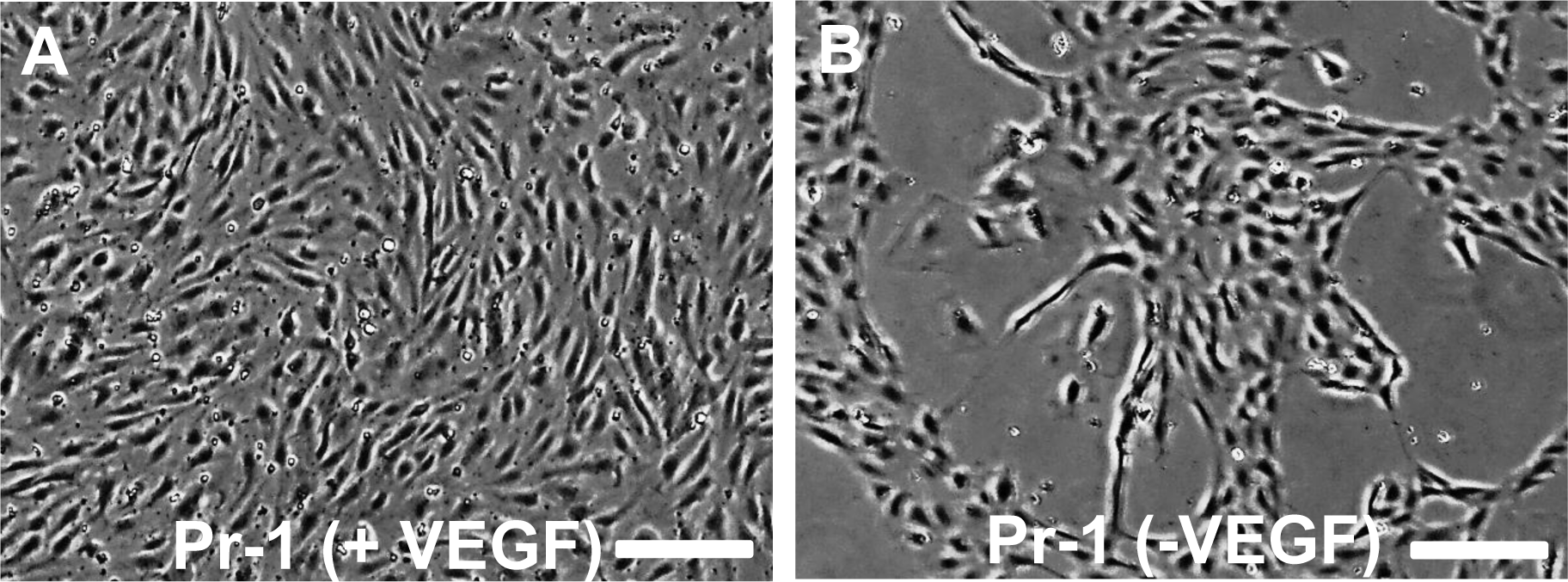
Cells differentiated according to the Pr-1 protocol require VEGF to form and maintain uniform monolayers. Bright-field micrographs illustrating the effect of a VEGF deprivation on a monolayer of Pr-1 derived cells. The cells were produced and amplified in the presence of VEGF according to the Pr-1 protocol until the formation of a uniform monolayer **(A)** and then cultured overnight in the same medium without VEGF **(B)**. Scale bars: 0.2 mm.

**Supplementary Figure 2.**
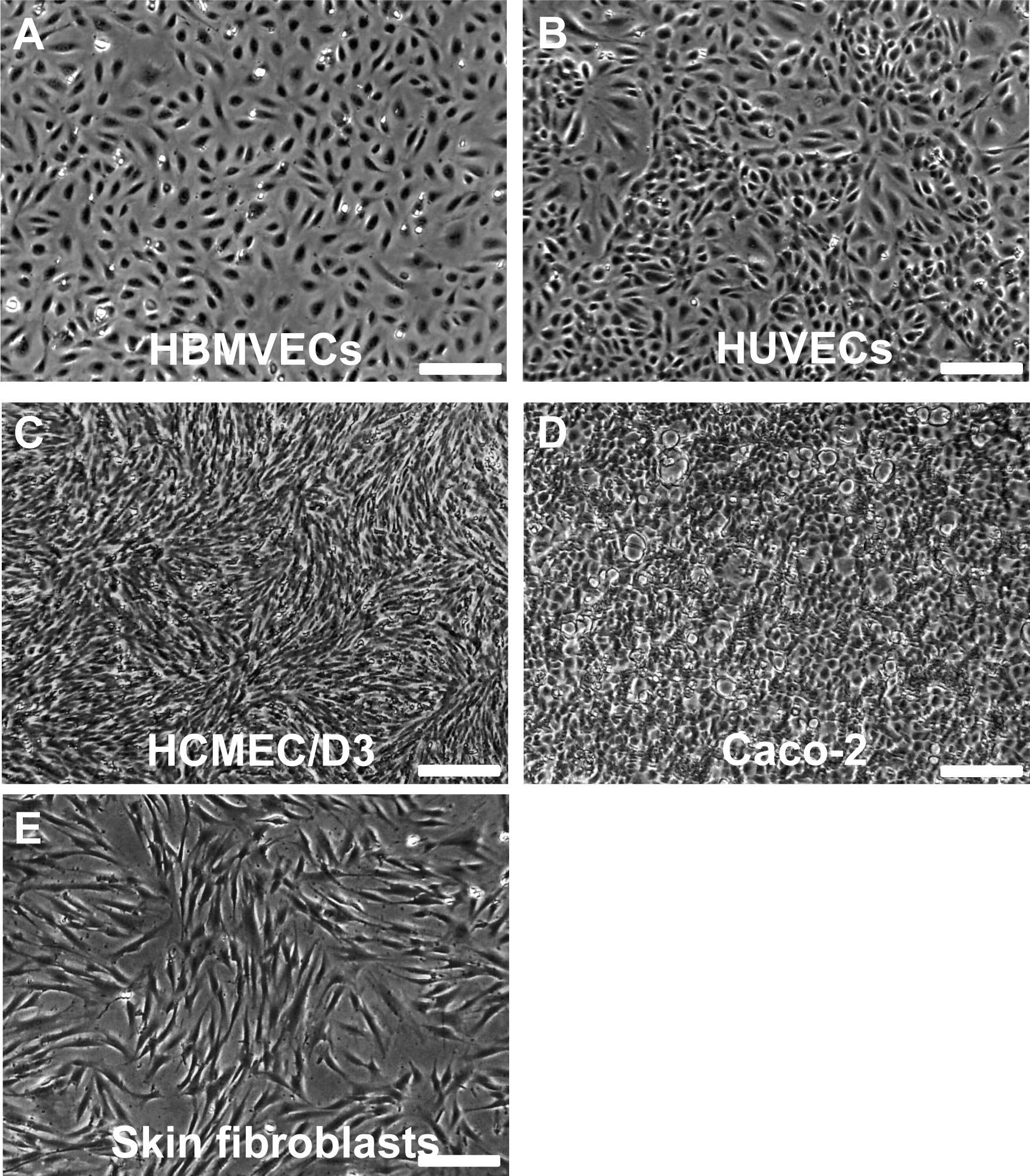
Representative morphologies of cell lines and primary cells used as controls throughout the study. Bright-field micrographs illustrating the morphology of the cells used as control in the present study. Scale bars: 0.2 mm.

**Supplementary Figure 3.**
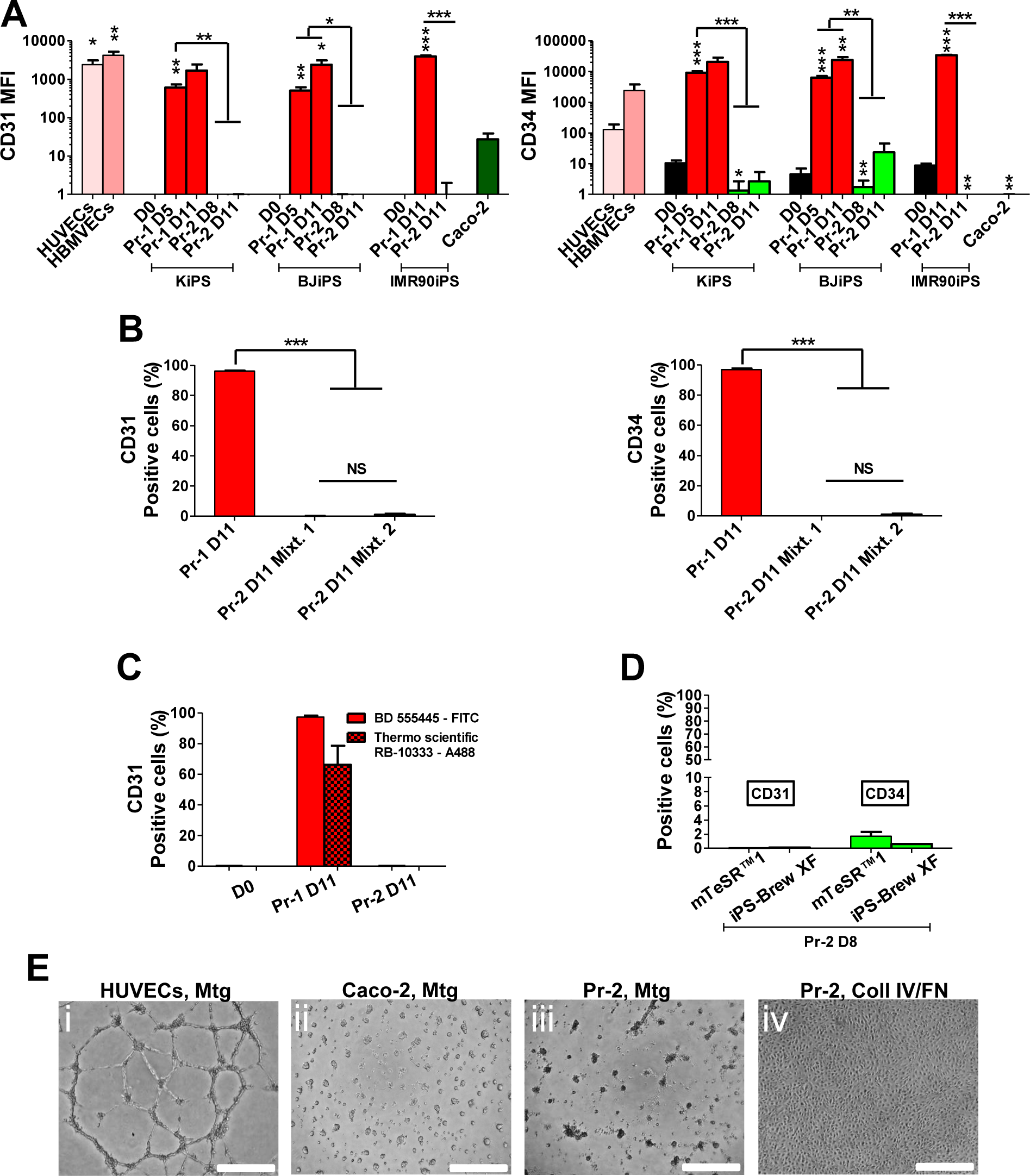
Evaluation of the endothelial phenotype in iPSC-derived cells obtained with Pr-1 or Pr-2 differentiating protocols. A) The expression of CD31 and CD34, two EC markers, was assessed by flow cytometry and quantified as mean fluorescence intensity (MFI). The bar graphs display results obtained from three different hiPSC lines with Pr-1 and Pr-2, at different stages of the two protocols as indicated. EC (HUVECs, HBMVECs) and EpC (Caco-2) cells were respectively used as positive and negative controls in these experiments. B) Bar graphs showing the percentage of CD31 or CD34 positive cells, as assessed by flow cytometry, in KiPS-derivatives obtained with Pr-2. Pr-2-derived cells were either purified on plates coated with a mixture of collagen type IV/ fibronectin at 10 µg/mL (Mixt. 1) as previously described (Molino et al., 2014) or with a mixture of 400 µg/ml collagen type IV and 100 µg/ml fibronectin (*i.e.* 200 and 50 µg/cm^2^, respectively) (Mixt. 2) as previously reported (Lippmann et al., 2012). Pr-1-derived cells were used as control. C) Assessment of CD31 expression by flow cytometry on fixed/permeabilized KiPS-derivatives obtained with Pr-2. Herein, two different anti-CD31 antibodies were tested: the antibody used in the present study (BD 555445) was compared with the one (RB-10333) used by Lippmann *et al*. (Lippmann et al., 2012). Undifferentiated KiPS and differentiated Pr-1-derived cells were used as negative and positive controls, respectively. D) Bar graph displaying the number of CD31 or CD34 positive cells obtained from unpurified BJiPS-derivatives obtained with Pr-2 (D8) and assessed by flow cytometry. Of note, to avoid confounding factor due to the iPSC medium, cells obtained from undifferentiated hiPSCs maintained in the medium used in the present study (iPS-BREW XF) were compared to those obtained from hiPSCs maintained in the medium used by the original report on Pr-2 (mTeSR^TM^1) (Lippmann et al., 2012). E) Representative bright-field micrographs of the indicated cells 24h after Matrigel capillary-like tube formation assay (panels i, ii and iii). On the last panel (iv), Pr-2-derived cells from the same cell suspension as the one used for panel iii were seeded onto collagen type IV / fibronectin coated plates to test the cell viability. Scale bar: 500 µm. Values reported are the mean (± SEM) of at least three independent differentiations/cultures with *p ≤ 0.05, **p ≤ 0.01, and ***p ≤ 0.001, using Student’s t-test. NS: not significant.

**Supplementary Figure 4.**
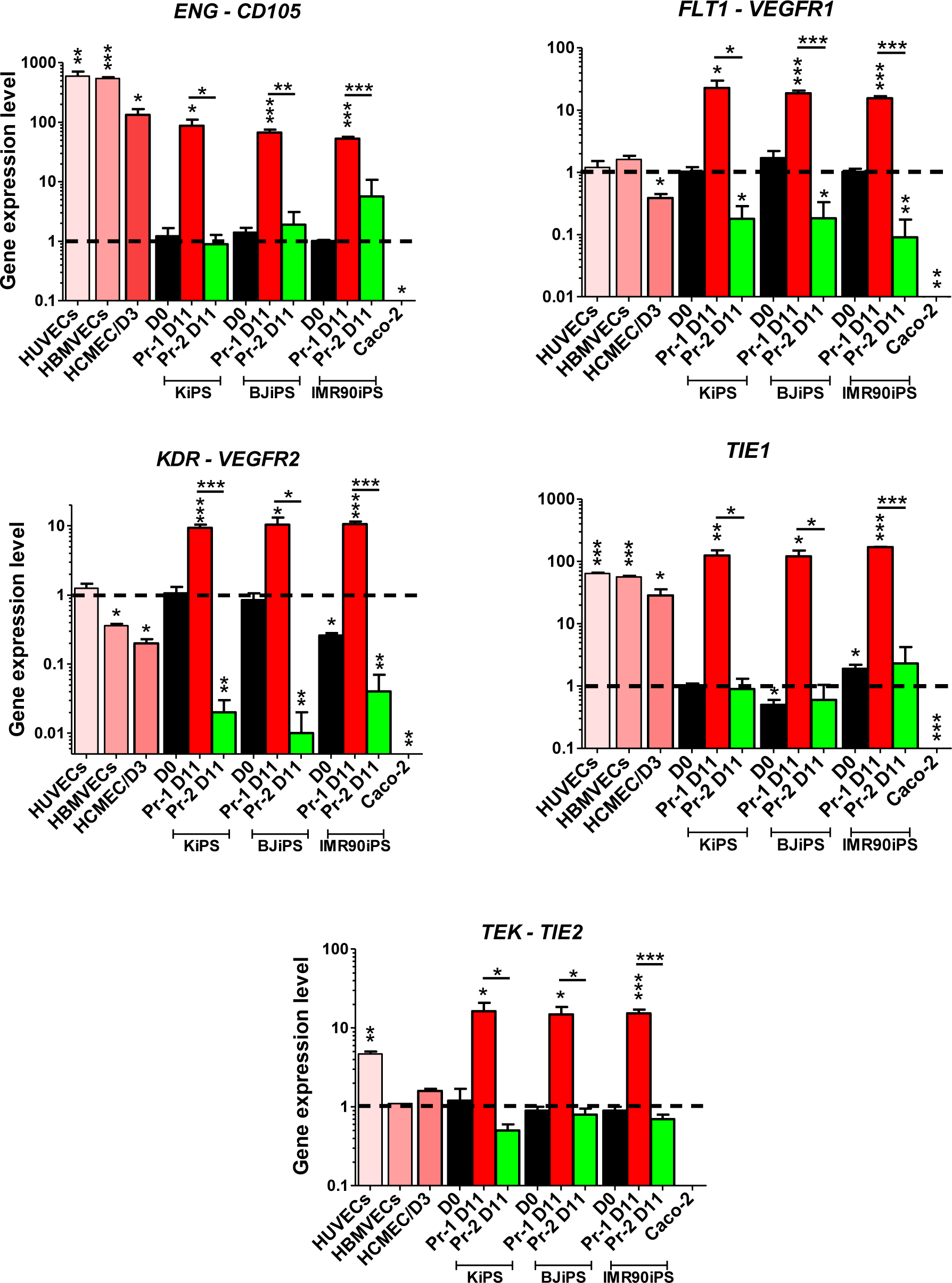
Comparative expression of a set of EC markers between Pr-1- and Pr-2-derived cells. Bar graphs showing the mRNA expression profile of a set of 5 EC markers, as indicated, in hiPSC-derivatives obtained with both protocols, Pr-1 and Pr-2. EC (HUVECs, HBMVECs, HCMEC/D3) and EpC (Caco-2) cells were used as controls. Values reported are shown as fold change relative to the value of their respective undifferentiated hiPSCs (D0) and are represented as a mean (± SEM) of at least three independent differentiations/cultures with *p ≤ 0.05, **p ≤ 0.01, and ***p ≤ 0.001, using Student’s t-test.

**Supplementary Figure 5.**
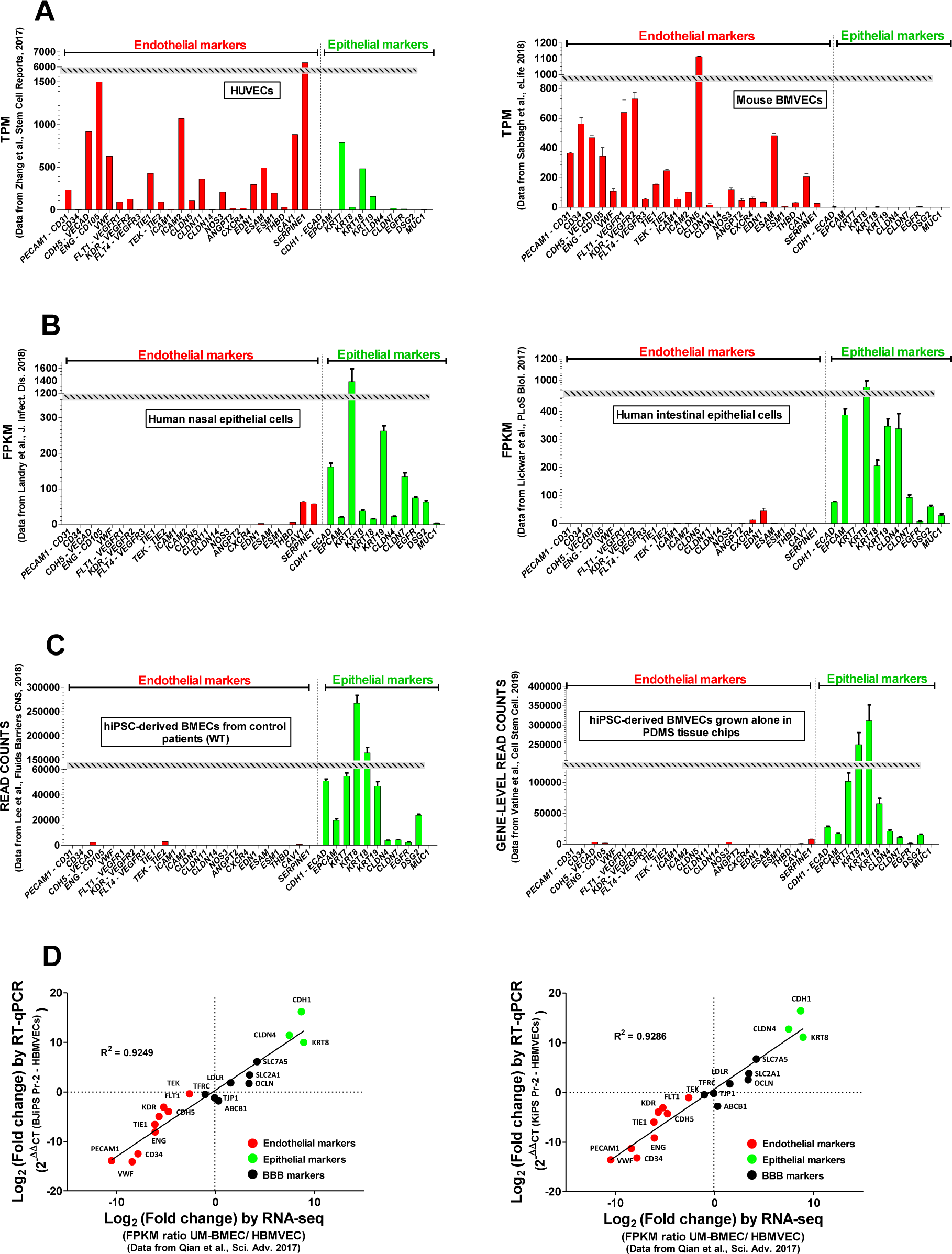
Transcriptomic profiling from Pr-1 and Pr-2 derivatives reveal two distinct phenotypes. RNAseq data sets were extracted from the GEO database in order to obtain the gene expression profile of a selected set of EC and EpC markers according to previous studies (A-C). **A)** Gene expression profile of primary ECs: HUVECs (accession number: GSE93511, (Zhang et al., 2017)) and mouse BMVECs (accession number: GSE111839 (Sabbagh et al., 2018)). **B)** Gene expression profile of human primary EpCs: nasal EpCs (accession number: GSE 107898 (Landry and Foxman, 2018)) and intestinal EpCs (accession number: GSE94935 (Lickwar et al., 2017)). **C)** Gene expression profile of hiPCS-derived cells differentiated according to “Pr-2-like” protocols (accession numbers: GSE108012 and GSE129290 (Lee et al., 2018, Vatine et al., 2019). **D)** Correlative analysis between the relative gene expression of the selected sets of EC, EpC and BBB markers as measured in the present study by RT-qPCR and in a previous study by RNAseq (Qian et al., 2017). The value represents the log2 of the fold change between HBMVECs and BJiPS or KiPS-derivatives from the Pr-2 protocol (RT-qPCR) or a “Pr-2-like” protocol (UM-BMEC, RNAseq). TPM: Transcripts Per Kilobase Million.

**Supplementary Figure 6.**
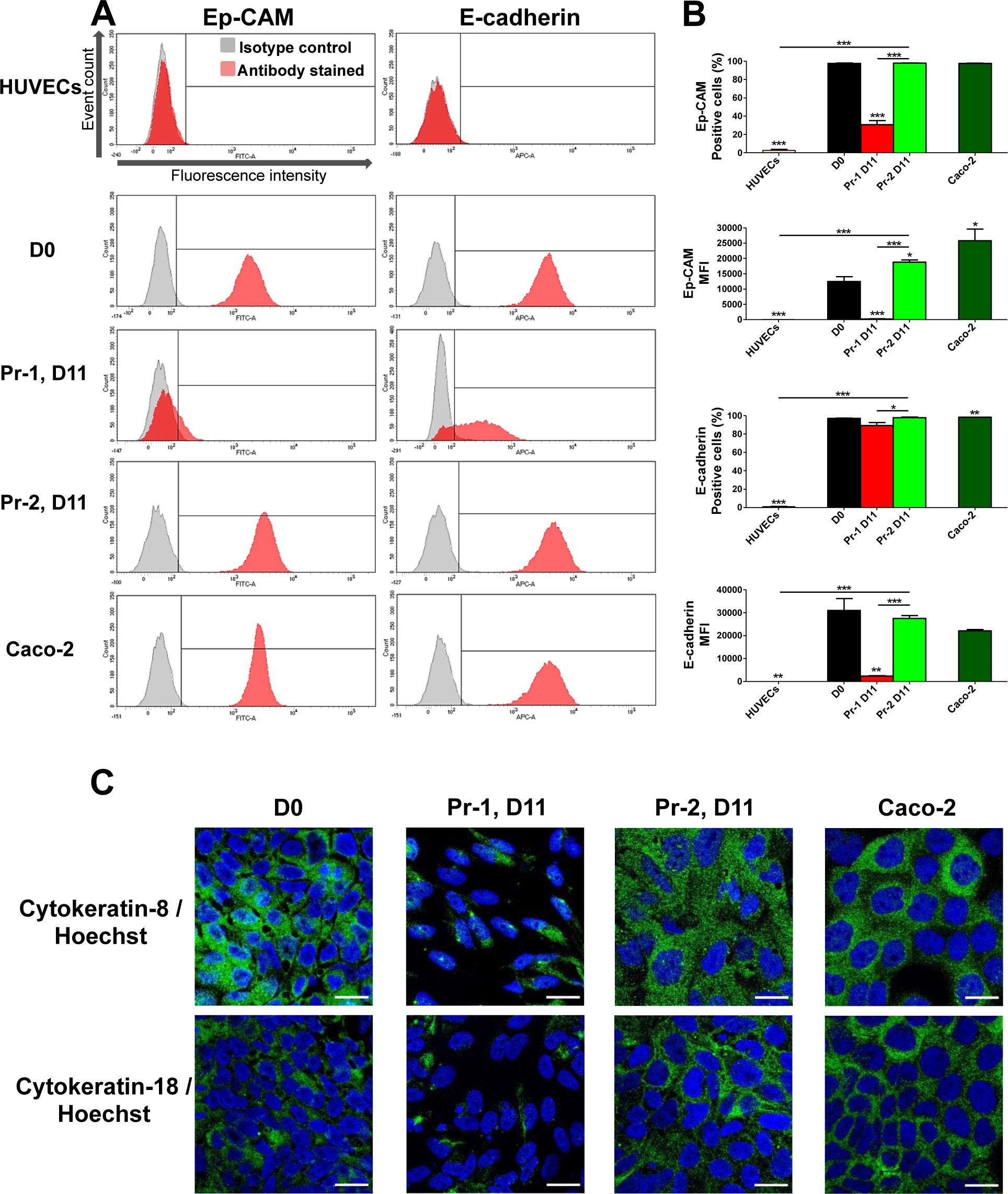
Pr-2-derived cells express typical epithelial markers. **A)** Representative flow cytometry results showing fluorescence intensity and cell count number of stained cells for two EpC markers (Ep-CAM and E-cadherin) in differentiated cells according to Pr-1 or Pr-2 protocols, at day 11 (D11) of differentiation. Appropriate isotype controls (grey plot) were used throughout the study. HUVECs and Caco-2 were used as EC and EpC positive controls, respectively. Undifferentiated iPSCs (KiPS) were also analyzed. The analyses revealed that Pr-2-derived cells strongly expressed EpCs markers but were barely detectable (or at low intensity) in Pr-1-derived cells. **B)** Bar graphs showing the percentage of positive cells for the indicated EpC markers as well as their mean fluorescent intensity (MFI) as measured by flow cytometry. Data are represented as the mean (± SEM) of at least three independent differentiations/cultures with *p ≤ 0.05, **p ≤ 0.01, and ***p ≤ 0.001, using Student’s t-test. **C)** Representative fluorescent micrographs showing immunostainings against two EpC markers (Cytokeratin-8 and Cytokeratin-18, green). Nuclei were counterstained with Hoechst (blue). Scale bar: 20 µm.

**Supplementary table 1:** List of cell lines and primary cells used in this study.

| Cell line | Cell line aliases | Description | Supplier / Origin | Passage |
| --- | --- | --- | --- | --- |
| **KiPS** | **KIPS 4F2** | Human iPSCs reprogrammed from juvenile foreskin keratinocytes by retroviral transduction (*OCT4*, *SOX2*, *KLF4*, *c-MYC*) (Aasen et al., 2008) | Gene Expression Laboratory, Salk Institute for Biological Studies | 43 - 65 |
| **BJiPS** | **Fibro-Epi** | Human iPSCs reprogrammed from neonatal foreskin fibroblast by episomal transfection ( *OCT3/4*, *shRNA-p53*, *SOX2*, *KLF4*, *LMYC* and *LIN28*) (Xia et al., 2012) | Gene Expression Laboratory, Salk Institute for Biological Studies | 35 - 55 |
| **IMR90iPS** | **iPS(IMR90)-4** | Human iPSCs reprogrammed from fetal fibroblasts by lentiviral transduction (*OCT4*, *SOX2*, *NANOG*, *LIN28*) (Yu et al., 2007) | WiCell Research Institute | 51 – 65 |
| **HUVECs** |  | Primary human umbilical vein endothelial cells from pooled donors | Promocell | 5 - 8 |
| **Caco-2** | **ATCC® HTB-37™** | Human epithelial colorectal adenocarcinoma cells | American Type Culture Collection (ATCC) |  |
| **HBMVECs** |  | Primary human brain microvascular endothelial cells | Angio-Proteomie / Pelobiotech | 3-5 |
| **HCMEC/D3** |  | Immortalized human cerebral microvascular endothelial cell line | Laboratory of Dr. P-O Couraud, Institut Cochin | 30 - 35 |
| **Skin fibroblasts** | **GM03348** | Human primary dermal fibroblasts isolated from skin biopsy | Coriell Institute | 12 - 15 |

**Supplementary table 2:**
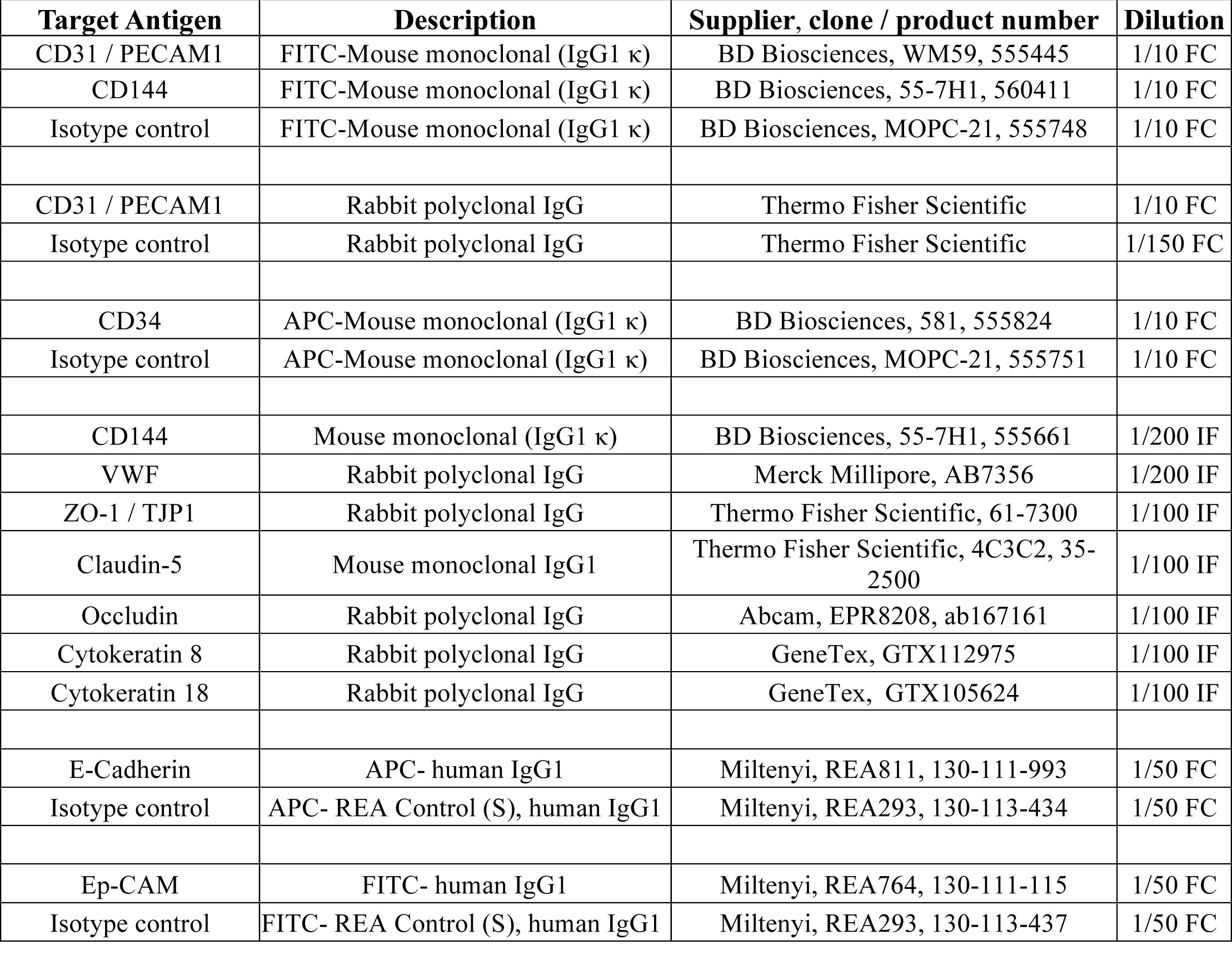
Primary antibodies list.

**Supplementary table 3:**
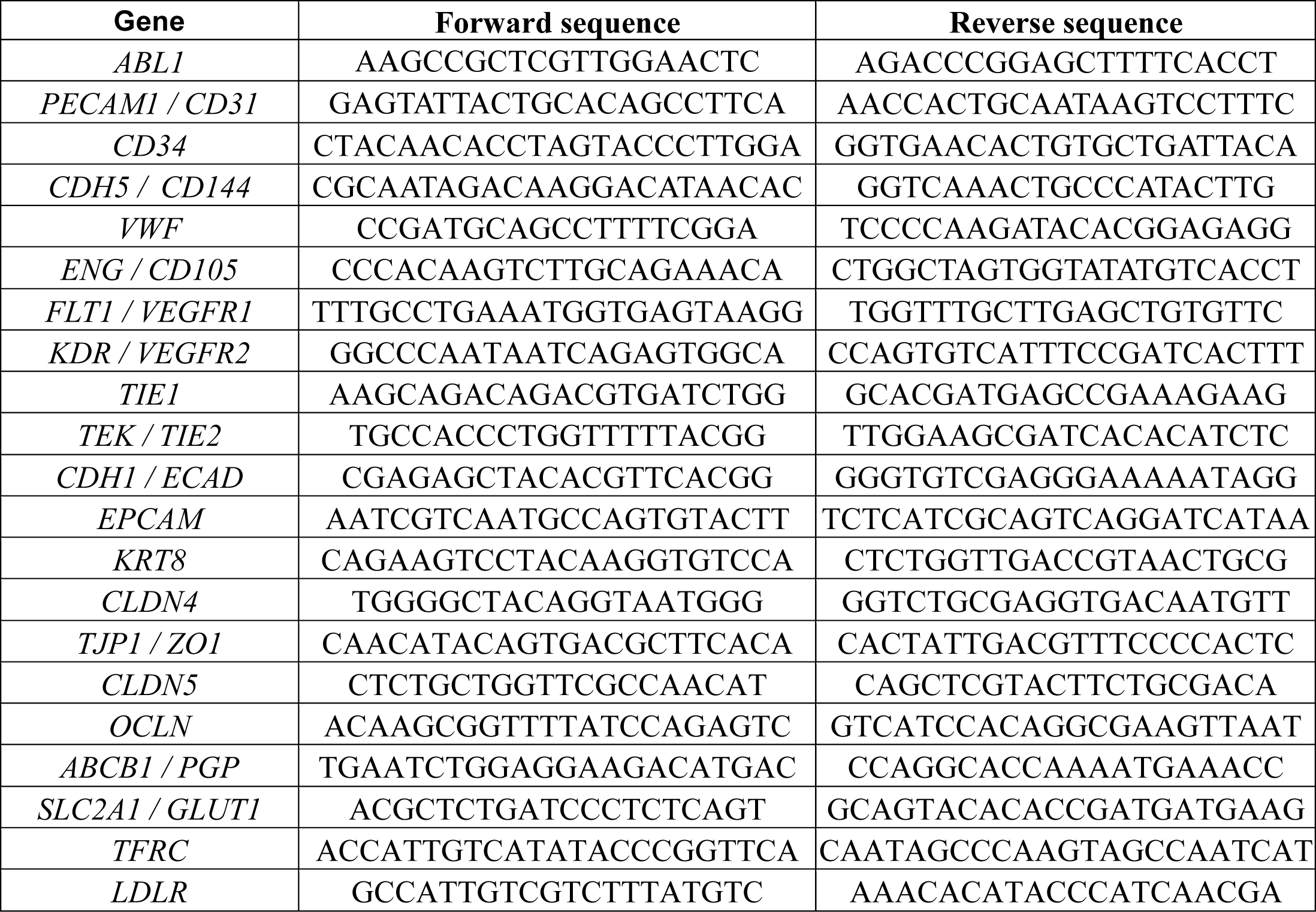
Primers used for RT-qPCR experiments.

